# Transcriptional response to chronic long-access fentanyl self-administration in rat habenula and amygdala

**DOI:** 10.1101/2025.11.25.690517

**Authors:** Robin Magnard, Daianna Gonzalez-Padilla, Ege A. Yalcinbas, Emma Chaloux-Pinette, Nicholas J. Eagles, Michael S. Totty, Patricia H. Janak, Leonardo Collado-Torres, Kristen R. Maynard

## Abstract

Fentanyl is a potent synthetic opioid associated with overdose. However, little is known about fentanyl-induced molecular adaptations in the habenula and amygdala, two brain regions implicated in opioid use and withdrawal. We performed bulk RNA-sequencing in the rat habenula and amygdala to identify transcriptomic changes associated with fentanyl intake. Rats self-administered intravenous saline or fentanyl over 22-24 days. Ninety minutes following the final session, brains were collected for transcriptomic profiling. In Hb, we identified 453 differentially expressed genes (DEGs) between saline and fentanyl rats, with upregulated genes associated with synaptic transmission and ionic conductance. In amygdala, we identified 3,041 fentanyl-associated DEGs with upregulated genes implicated in metabolic and vesicular functions. Downregulated genes in both regions were enriched for extracellular matrix functions. Integration of DEGs with single-cell RNA-sequencing data from rodents and humans revealed that fentanyl DEGs were enriched in specific habenula and amygdala cell type markers. Furthermore, fentanyl downregulated DEGs in amygdala were enriched in genes associated with risk for substance use disorders. Together, we define how fentanyl intake alters transcriptional programs in the rat habenula and amygdala, and we link these changes to specific human cell types and risk genes for neuropsychiatric disorders and addiction.

## Introduction

Over the past decade, overdose deaths involving opioids have been rising due to the prevalence of illicit use of synthetic opioids [1–3]. Fentanyl is a synthetic opioid 50-100 times more potent than other μ-opioid receptor (MOR) agonists, such as morphine and oxycodone [4], and it has disproportionately contributed to the rise in opioid-related deaths [2,5,6]. Beyond its potency, additional pharmacological properties that distinguish fentanyl from other MOR agonists include higher lipid solubility allowing more rapid cell membrane penetration and receptor binding [4]. These pharmacological differences have downstream molecular consequences, such as reduced sensitivity to reversal by the MOR antagonist, naloxone, compared to morphine [2,7]. Thus, a deeper understanding of the molecular adaptations induced by fentanyl intake is needed to identify pathways that can be targeted to mitigate its effects.

Opioids cause long-term molecular changes in the brain, especially reward circuits involving the habenula (Hb) and amygdala (Amyg) [8–10]. The Hb consists primarily of glutamatergic neurons, and acts as an “anti-reward” center encoding aversive states by modulating monoaminergic hubs such as the ventral tegmental area (VTA), dorsal raphe nucleus (DRN), and locus coeruleus (LC) [11,12]. The Hb has a high density of MOR [13], especially at the transition between the medial and lateral Hb divisions [14,15]. Optogenetic excitation of MOR-expressing Hb neurons promotes aversive states and elicits avoidance [16]. Hb MOR-expressing neurons have also been shown to mediate the aversive effects of naloxone-precipitated withdrawal [17]. While it is clear that opioid signaling in the Hb mediates distinct functional and behavioral responses, no one has yet investigated how opioids, especially potent synthetic opioids, such as fentanyl, alter the transcriptional landscape of this key hub of reward circuitry.

The Amyg includes several nuclei among which the basolateral amygdala (BLA) and central amygdala (CeA) play an important role in processing emotional valence and affective behaviors [18,19]. The BLA assigns value to environmental stimuli and is critical for associating the negative emotional state experienced during opioid withdrawal with discrete contexts and cues [20–22]. BLA also supports context dependent memories. Song et al., 2019 [23] showed that retrieving morphine-associated withdrawal memories requires BLA plasticity and coordinated activity within a BLA→prelimbic cortex loop. More recently, chronic oral fentanyl consumption and withdrawal were found to impair fear extinction learning and induce persistent hyperexcitability in BLA glutamatergic principal neurons [24]. In contrast, the CeA has been implicated in the negative affective state of opioid withdrawal [18,25]. Lesioning the central nucleus of the amygdala (CeA) blocks morphine withdrawal-induced conditioned place aversion [26]. A population of MOR-expressing neurons in the CeA is activated during fentanyl withdrawal and drives aversive symptoms and negative reinforcement [27]. Transient hyperactivity of this MOR-expressing CeA neuron population is tied to naloxone-precipitated withdrawal in mice subjected to repeated fentanyl injections compared to drug naive mice [27]. Furthermore, opioid-induced hyperalgesia can be produced by repeated fentanyl administration in rodents, and the CeA has been implicated in this aversive experience, which is also associated with chronic fentanyl self-administration [28,29]. While there is mounting evidence for the role of the Amyg in withdrawal and negative states following chronic fentanyl, the molecular adaptations that occur in this region during the trajectory of chronic fentanyl SA remain unexplored.

In rodents, acute or repeated injection of opioids induces gene expression dysregulation in the brain [30–34]. Similarly, chronic self-administration of opioids (i.e. oxycodone, morphine, and heroin) causes widespread molecular changes in reward circuitry nodes, including MOR-rich brain regions like the nucleus accumbens (NAc), dorsal striatum, ventral tegmental area (VTA), prefrontal cortex, and basolateral Amyg, with many of these changes occurring dynamically in different cell types across phases of use (i.e. intake, abstinence, withdrawal, and reinstatement) [9,35–41]. Recent reports have also characterized how chronic long-access fentanyl self-administration alters gene expression in reward circuits [42–44], but these studies did not include the Hb or Amyg. In postmortem human brain, limited studies have identified transcriptomic changes associated with OUD in the dorsal striatum, NAc, and dorsolateral prefrontal cortex [45,46]. However, given that many individuals with OUD engage in polysubstance use [47–49], it is difficult to identify fentanyl-specific transcriptional effects in postmortem human brains, reinforcing the utility of single substance self-administration models in rodents. In induced pluripotent stem cell (iPSC)-derived models, single-cell sequencing of forebrain organoids from patients with OUD showed that repeated exposure to different classes of opioids leads to unique transcriptional responses [50]. Together, these studies highlight the importance of understanding fentanyl-specific transcriptional programs across rodent and human Hb and Amyg.

Here, we performed bulk RNA-sequencing (RNA-seq) in the rat Hb and Amyg following chronic long-access fentanyl self-administration to identify shared and unique molecular alterations in these regions associated with fentanyl intake. In each region, we identified differentially expressed genes (DEGs) between saline and fentanyl rats and evaluated enrichment of these DEGs in cellular processes and molecular pathways. We integrated these data with publicly available single-nucleus (sn) and single-cell (sc) RNA-seq data from rodent and human Hb and Amyg [51–54] to explore whether particular cell types are associated with chronic fentanyl self-administration. Finally, using GWAS summary statistics, we investigated whether Hb and Amyg DEGs for chronic fentanyl self-administration are associated with higher risk for neuropsychiatric disorders, including OUD and other substance use disorders (SUD) [55–60].

## Materials and Methods summary

### Rats

Adult male Sprague-Dawley rats (ENVIGO, Frederick, MD) were used in all experiments (n = 8 fentanyl, n = 11 saline). Rats were single-housed in a temperature and humidity-controlled vivarium on a normal light cycle (12/12 hr. light/dark cycle, lights ON at 7 am) with ad libitum access to food and water. The protocol was approved by the Animal Care and Use Committee of Johns Hopkins University and was conducted in accordance with the National Institutes of Health Guidelines for the Care and Use of Laboratory Animals.

### Drugs

Fentanyl Citrate (Cayman Chemical) was diluted in 0.9% sterile saline at 10 ug/mL. Brevital sodium (Henry Schein) was diluted to 10 mg/mL with 0.9% sterile saline.

### Surgeries

Adult male rats, weighing 250-300 grams at the time of surgery, were anesthetized with isoflurane (3-5% induction, 1-2% maintenance) and administered the analgesic rimadyl (5 mg/kg, SC, Zoetis) and the antibiotic cefazolin (70 mg/kg, SC, West-Ward) prior to catheter implantation. A silastic jugular catheter, constructed as described previously [61], was threaded under the skin from a subcutaneous base in the mid-scapular region and inserted into the right jugular vein [61]. Catheters were flushed daily with ∼0.1 mL of sterile saline solution containing 10 mg/mL gentamicin sulfate (VetOne) and 100 IU/mL Heparin (Sagent). Once behavioral training began, catheters were flushed before and after self-administration (and daily on rest days), and were assessed daily for blood return. In the event that blood return was absent, rats were administered 0.1 mL of brevital sodium (10 mg/mL in sterile saline, Henry Schein). If the rat did not become ataxic within 10 seconds, the catheter was considered not patent and the rat was removed from the study.

### Self-administration training

After recovery, rats were trained in standard Med Associates operant chambers housed inside sound-attenuating chambers (Med Associates, St Albans, VT, USA). Operant chambers were fitted with two retractable levers (active and inactive), stimulus lights above each lever, speakers, and a house light. Boxes were controlled using Med-PC IV software (Med Associates, St Albans, VT, USA).

Rats were trained to lever-press on a fixed-ratio 1 schedule, initially in short-access 2-hour sessions, and then progressing to long-access 6-hour sessions (6 days per week). Both levers were extended for the entire session duration. The stimulus light above the active lever indicated drug (or saline) availability. For fentanyl rats, active lever presses resulted in infusion of 0.5 µg fentanyl dissolved in 50 µl sterile saline delivered over 2.8 seconds [62], while saline rats received an equivalent volume of sterile saline. Infusion was accompanied by a 2.8 second tone, followed by an additional 20 second timeout during which the stimulus light and house light were extinguished; active lever responses during this 22.8 sec interval were not reinforced. After 20 seconds elapsed, the house light and the stimulus light were re-illuminated to indicate availability of drug (or saline). Training occurred for 22-24 sessions, after which the brains were collected. Behavioral data for all the rats included in this study are provided (**Figure 1**, **Figure S1**, **Table S1, Table S2**, **Table S3**). The escalation ratio, used as an additional metric of infusion escalation across sessions, was calculated by normalizing each rat’s LgA infusion counts relative to its infusion count on the first LgA session.

**Fig. 1.**
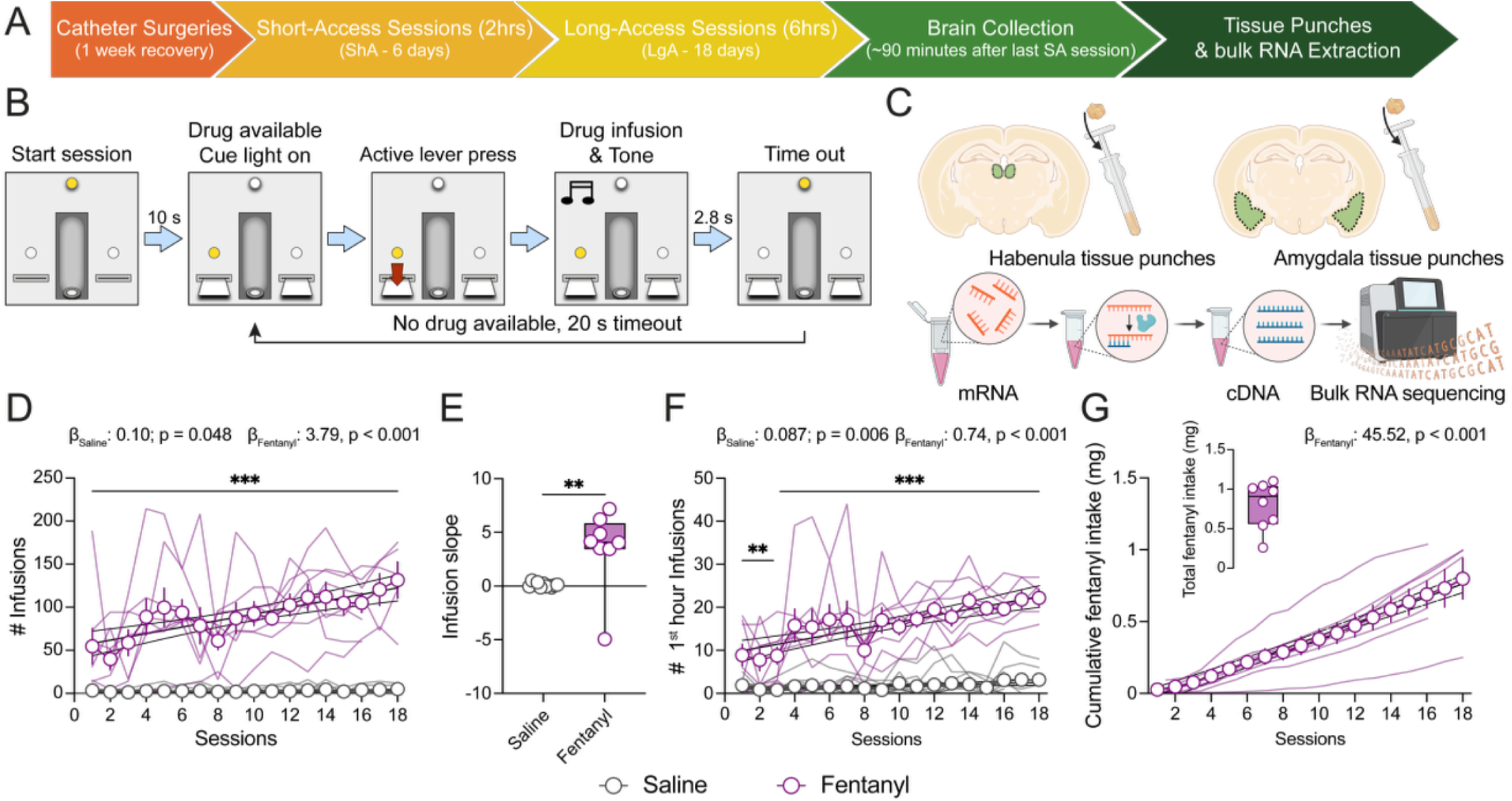
Escalation of fentanyl intake during chronic long-access intravenous self-administration. (**A**) Experimental timeline. Rats were implanted with jugular catheters for intravenous delivery of either saline or fentanyl. Rats first underwent six 2-hour acquisition sessions (Short-Access, ShA), followed by 16-18 daily 6-hour sessions (Long-Access, LgA). (**B**) Intravenous self-administration task design. Fentanyl or saline infusion was contingent upon active lever press on a fixed-ratio 1 (FR1) schedule of reinforcement. Drug or saline availability was signaled by cue light illumination and extinction of the house light. (**C**) Schematic of bilateral habenula and amygdala tissue collection for bulk RNA sequencing. (**D**) Mean number of infusions per LgA session for saline and fentanyl rats (two-way RM ANOVA, substance effect: F_1,17_ = 79.38, p < 0.001; session effect: F_17,273_ = 5.98, p < 0.001; substance x session interaction: F_17,273_ = 5.42, p < 0.001; Student Newman-Keuls post hoc pairwise comparison; ANCOVA substance slope difference: F_1,322_ = 32.07, p < 0.001). (**E**) Individual rat infusion slopes across the LgA sessions (t-test, substance effect t_17_ = 3.14, p = 0.006). (**F**) Mean number of infusions occurring within the first hour of each LgA session for saline and fentanyl rats (two-way RM ANOVA, substance effect: F_1,17_ = 73.59, p < 0.001; session effect: F_17,273_ = 7.93, p < 0.001; substance x session interaction: F_17,273_ = 5.59, p < 0.001; Student Newman-Keuls post hoc pairwise comparison; ANCOVA substance slope difference: F_1,322_ = 32.76, p < 0.001). (**G**) Mean cumulative fentanyl intake (µg) across LgA sessions. Insert: total overall amount of fentanyl intake per rat during ShA and LgA sessions. Data shown as mean across rats ± SEM, superimposed with individual rat data points. Black lines represent linear regression with 95% confidence intervals. Boxes extend from the 25^th^ to 75^th^ percentiles; lines within the boxes represent the median; whiskers indicate the minimum and maximum values. Saline: n = 11 rats; Fentanyl: n = 8 rats. * denotes p < 0.05; ** denotes p < 0.01; and *** denotes p < 0.001.

### Brain tissue extraction and bulk RNA-sequencing

Rats were lightly anesthetized with isoflurane and rapidly decapitated 60-90 minutes after the last long-access self-administration session, and extracted brains were fresh frozen in isopentane and stored at −80 °C. For tissue punching, brains were sectioned into ∼2 mm coronal slabs using a rat brain matrix (Stainless Steel Alto Coronal, 1.0 mm matrix, Small Rat 175-300gm), and bilateral tissue punches were collected from the Hb (#39443001RM, Leica Biosystems, nominal diameter 1.25 mm) and Amyg (nominal diameter 1.25 mm) on petri dishes placed in a dry ice bucket. Punched tissue was ejected into tubes sitting in dry ice, and sample tubes were stored at −80 °C until ready for RNA processing. Total RNA was isolated by using TRIzol Reagent homogenization (#15596018, Invitrogen) and chloroform layer separation. RNA was then purified using an RNeasy Micro (#74004, Qiagen) kit with an RNase-Free DNase step (Mat. No. 1023460, Qiagen) according to manufacturer’s instructions. RNA concentration and purity were measured using a NanoDrop Eight (ThermoScientific). RNA quality control assays were performed on the Agilent 2100 Bioanalyzer, and the RNA integrity number for all samples ranged from 7.2 to 8.6 (mean ±SEM = 7.9 ± 0.06).

Ribosomal RNA depletion and library preparation was performed including ERCC spike-ins. Specifically, Illumina Stranded Total RNA Prep with Ribo-Zero Plus Kit was used for library preparation from the habenula samples (10 ng of RNA input), and TruSeq Stranded Total RNA with Ribo-Zero Human/Mouse/Rat Gold Kit was used for library preparation from the amygdala samples (100 ng of RNA input). RNA-sequencing was carried out by Psomagen with the following configuration: ∼80M paired-end (PE) reads per sample on an Illumina NovaSeq 6000 S4 (152 bp PE).

### RNA-seq data processing and quality control analysis

Read quality control, read alignment, and gene expression quantification against the rat genome mRatBN7.2 (release 109) was performed using *SPEAQeasy* v87ba0b4 [63]. Lowly expressed genes were filtered and raw counts were transformed to counts per million in a log_2_-scale (log_2_-CPM), normalizing by sample library size and RNA composition [64]. Sample QC metrics (**Table S3, Table S4**) were examined across brain regions and substances (**Figure S2**), RNA extraction batches (**Figure S3**), and number of total self-administration sessions (**Figure S4**). Quality differed between brain regions, potentially due to the different Illumina kits used for library preparation, and therefore Hb and Amyg samples were analyzed separately in downstream exploratory analyses (**Figure S2**). Outlier identification of sample QC metrics (**Figure S5**) and Principal Component Analysis (**Figure S6**) revealed 4 Hb and 4 Amyg outlier samples (**Figure S7**), but manual inspection of all QC metrics considered for all these 8 samples supported their inclusion for downstream differential gene expression (DGE) analysis (**Figure S7**). A final dataset of 16,708 genes across 33 samples (8 fentanyl and 8 saline Hb samples, and 8 fentanyl and 9 saline Amyg samples) was used for DGE analysis.

### Differential Gene Expression (DGE)

DGE between fentanyl vs. saline self-administration and for behavioral traits among fentanyl-administered rats (i.e. the slope of fentanyl infusions across each LgA self-administration session 1^st^ hour, total fentanyl intake across LgA self-administration sessions, and last LgA self-administration session fentanyl intake), was assessed separately in Hb and Amyg using *limma*-*voom* v3.58.1 [65]. In each DGE model, gene expression was adjusted for batch effects and quality control metrics selected based on their correlations with other covariates and their contributions to gene expression variance (**Figure 2**, **Figure S8**). Genes with FDR-adjusted *p*-value<0.05 were considered differentially expressed genes (DEGs).

**Fig. 2.**
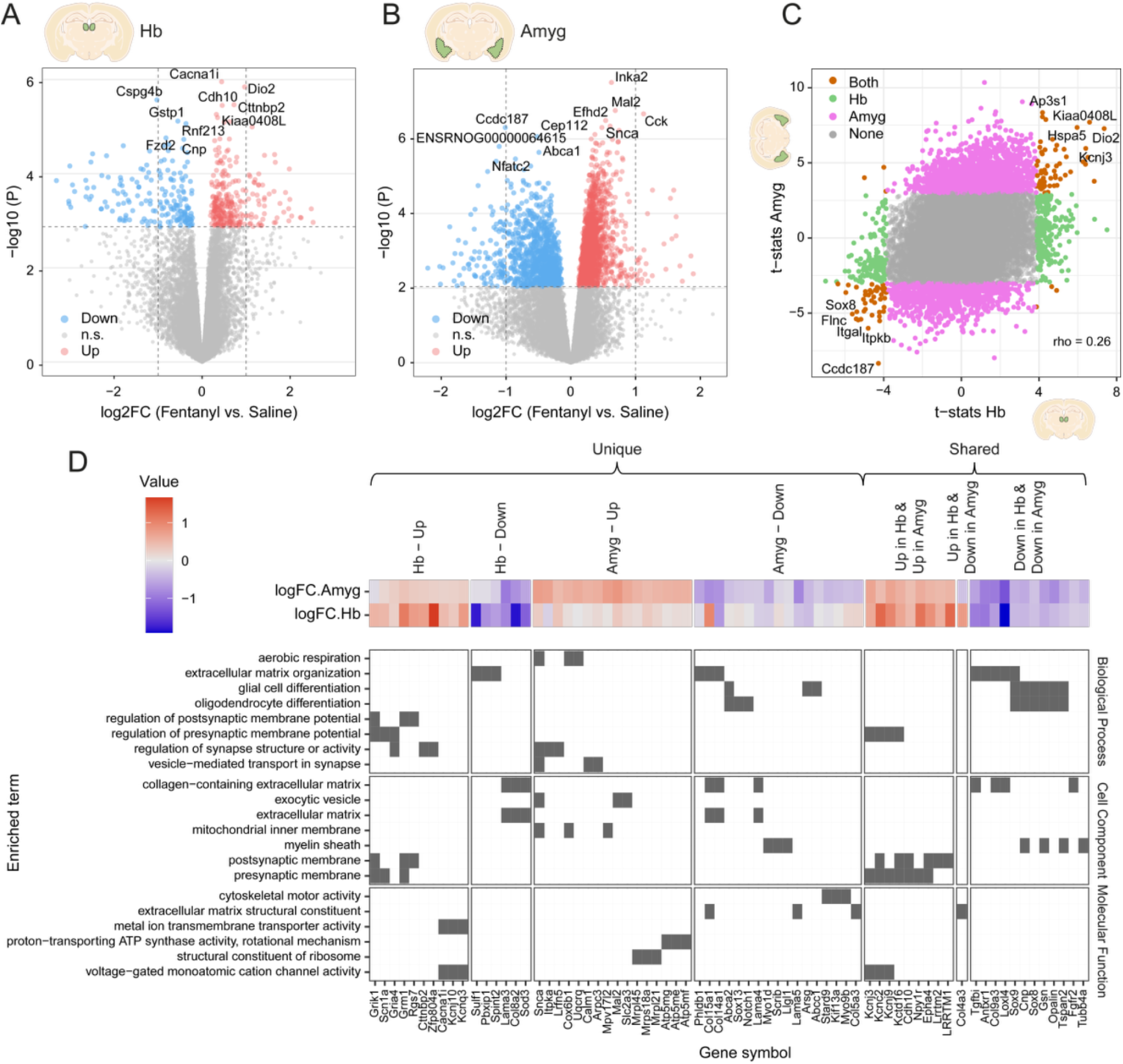
Transcriptional changes and gene ontology (GO) enrichment analysis in Hb and Amyg following chronic LgA fentanyl self-administration. (**A-B**) Volcano plots showing up- and down-regulated genes for fentanyl vs. saline in Hb (**A**) and Amyg (**B**). Horizontal dashed line demarcates significance threshold at FDR=0.05. Vertical dashed lines demarcate log_2_-fold change values of -1 and 1. The top five most significantly up- and down-regulated DEGs are labeled. (**C**) Substance (fentanyl vs. saline) t-statistic correlation plot in Amyg vs. Hb. Pink points are unique DEGs in Amyg. Green points are unique DEGs in Hb. Orange points are shared DEGs in Hb and Amyg. (**D**) Biological processes, cellular components, and molecular functions dysregulated by fentanyl in Hb and Amyg, as identified by Gene Ontology (GO) enrichment analysis of DEGs. Tile plot displays DEG (x-axis) membership to a significantly enriched GO term as a filled tile. Key DEGs per term are shown, categorized by their unique or shared up- and down-regulation in Hb and Amyg. Top heatmap shows DEG mean-centered log_2_FC in Hb and Amyg. Related to **Figure S9**, **Figure S10**, **Table S5**, **Table S6**, **Table S8**, **Table S9**.

### Functional enrichment analysis

Gene Ontology (GO) biological processes, molecular functions, and cellular components, and KEGG pathways affected by fentanyl self-administration in Hb and Amyg were found by over-representation analysis implemented in *clusterProfiler* v4.10.0 [66] (**Figure 2, Figure S10**). A total of 14,066 expressed genes, assessed for fentanyl vs. saline DGE, and with available EntrezIDs, were used as the background universe.

### Cell type enrichment analysis

The cell type specificity of DGE results in rat Hb and Amyg for fentanyl vs. saline self-administration was measured defining marker genes for fine and broad cell types in control rat Amyg [54], control human epithalamus [52] and Amyg [53], and control mouse Hb [51], with the *MeanRatio* method from *DeconvoBuddies* v0.99.0 [67]. Only cell types or clusters with at least 10 cells were considered. Rat orthologs of human and mouse cell type marker genes were obtained with *biomaRt* v2.56.1 [68]. The sets of cell type marker genes in rat were assessed for their over-representation among the rat DEGs in Hb and Amyg applying one-sided Fisher’s exact test (**Figure 3**, **Figure 4**). The set of genes assessed for DGE was taken as background (n = 16,708 genes).

**Fig. 3.**
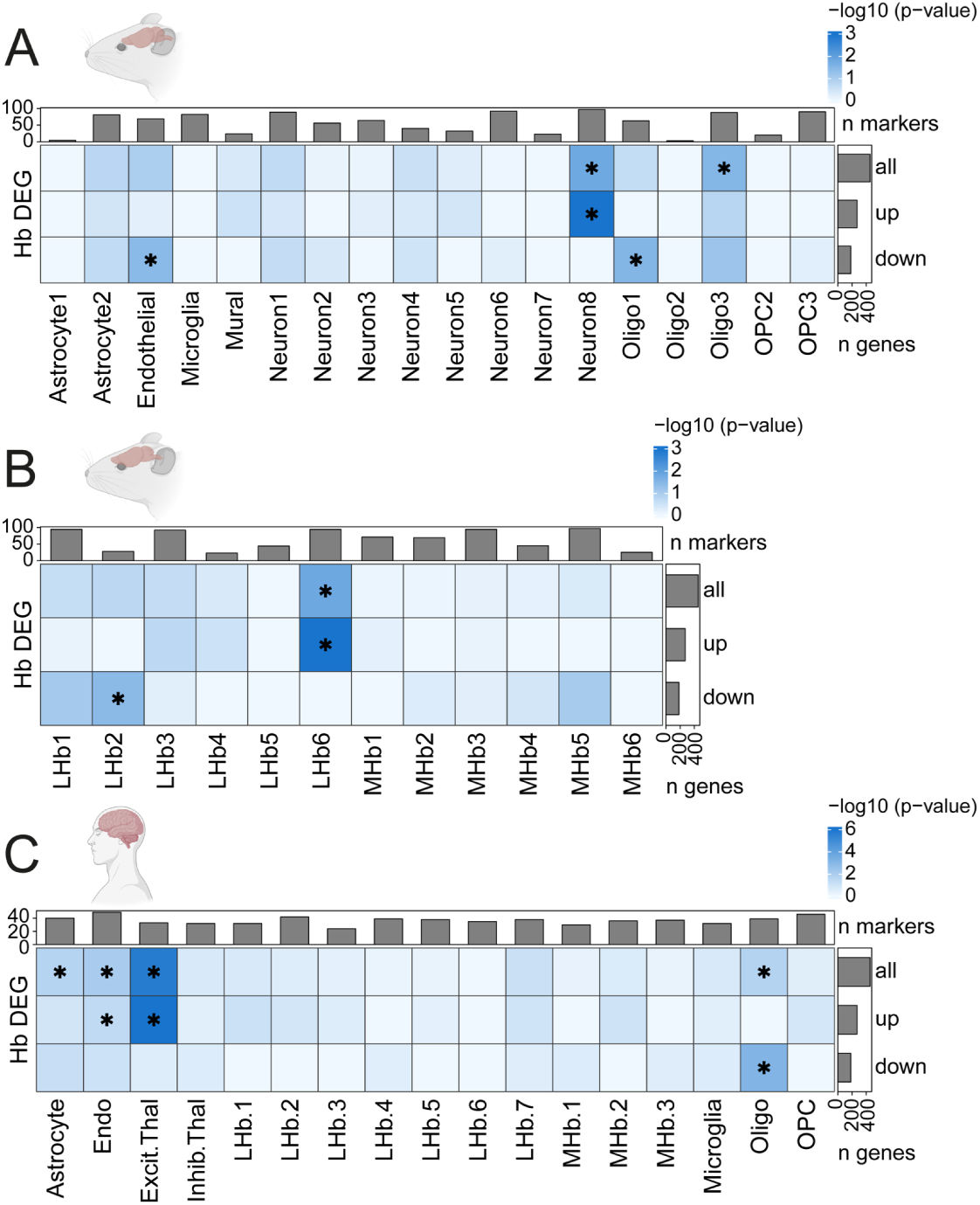
Enrichment of rat Hb fentanyl DEGs in specific human and rodent habenula cell types. (**A-B**) Enrichment of fentanyl vs. saline DEGs (FDR < 0.05) found in rat Hb (y-axis), among the rat orthologues of (**A**) the top 100 markers for broad-level cell type clusters and (**B**) top 100 markers for neuronal subtypes previously identified in mouse habenula by Hashikawa et al. [51] (**C**) Enrichment of rat fentanyl vs. saline DEGs among the rat orthologues of the top 50 markers for broad-level non-neuronal and fine-level neuronal subtypes we previously identified in human Hb/epithalamus in Yalcinbas et al. [52]. Note that mouse (Hashikawa et al. [51]), and human (Yalcinbas et al. [52]) Hb cell type names were not defined to match across datasets. Bar plots along the y-axis depict the number of Hb DEGs that were upregulated, downregulated, or all. Bar plots on top depict the number of rat orthologues used to define each cell cluster. The shade of blue indicates DEG enrichment p-value for a cell type in the -log_10_ scale. * denotes p < 0.05. Related to **Figure 2**, **Figure S11**, **Table S11**, **Table S12**.

**Fig. 4.**
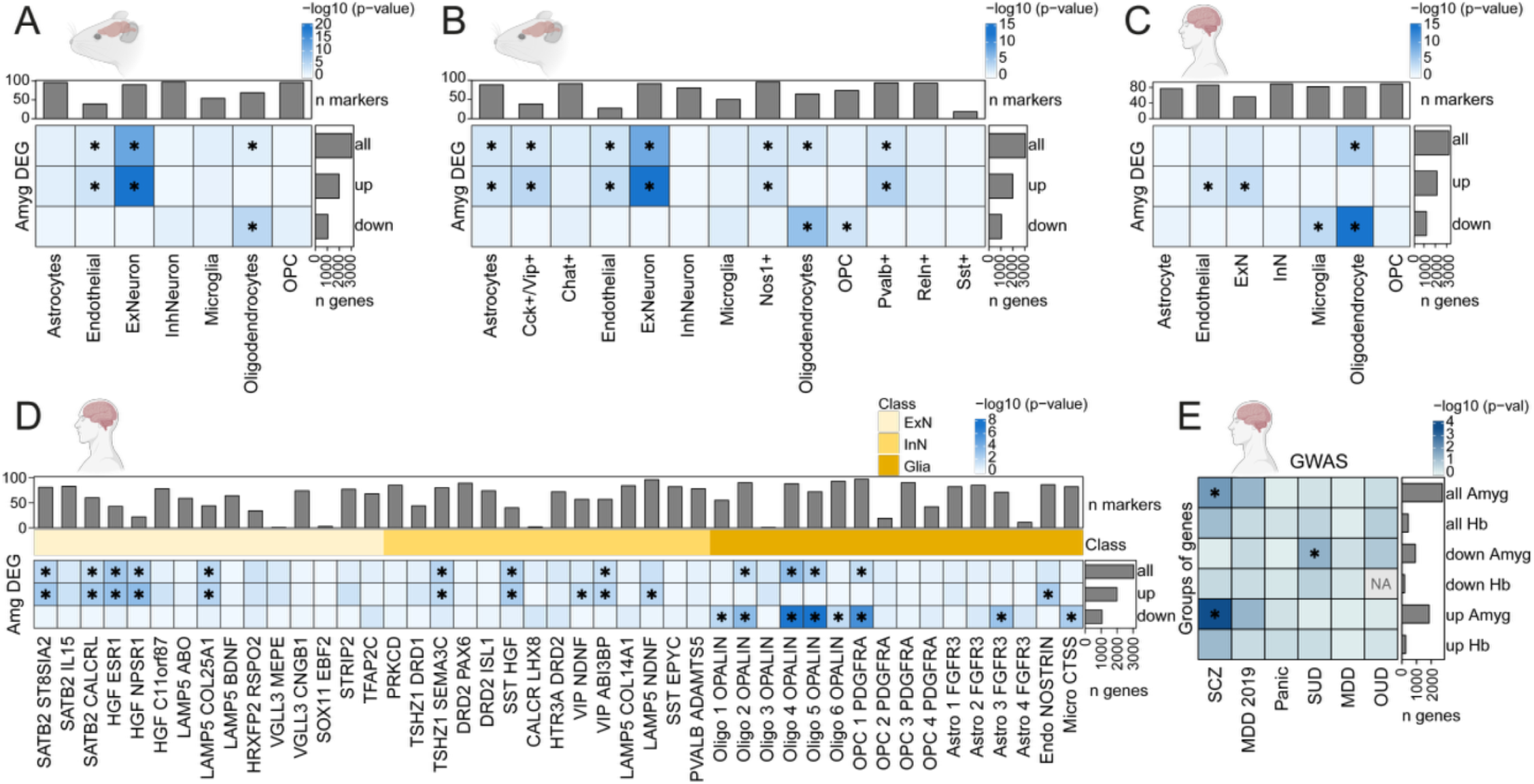
**Enrichment of rat Amyg fentanyl DEGs in specific human and rodent amygdala cell types and association of Hb and Amyg rat DEGs with psychiatric disorders**. Enrichment of fentanyl vs. saline DEGs (FDR < 0.05) found in rat Amyg (y-axis), among (**A**) the top 100 markers for broad-level and (**B**) fine-level cell type clusters identified in rat Amyg by Zhou et al. 2023 [54] among the rat orthologues of the top 100 markers for (**C**) broad-level and (**D**) fine-level cell types identified in the human Amyg by Yu et al. 2023 [53]. Bar plots along the y-axis depict the number of Amyg DEGs that were upregulated, downregulated, or all. Bar plots on top depict the number of marker genes used to define each cell cluster. The shade of blue indicates DEG enrichment p-value for a cell type in the -log_10_ scale. *p < 0.05. The light yellow, yellow, and orange colors indicate the cell type class: excitatory neuron (ExN), inhibitory neuron (InN), or glia. (**E**) *MAGMA* gene-set enrichment analysis of gene-level Genome-Wide Association Study (GWAS) signals in human orthologs of rat Hb and Amyg fentanyl vs. saline DEGs. The y-axis categorizes fentanyl vs. saline DEGs into upregulated, downregulated, or all, by brain region (Hb or Amyg). The x-axis represents the 5 GWAS datasets that were probed. The shade of blue indicates enrichment p-value in the -log_10_ scale. * denotes FDR-adjusted p < 0.05. Given only 1 human gene whose rat ortholog was downregulated in Hb had OUD-associated SNPs, *MAGMA* couldn’t perform enrichment for this gene set (returned NA). SCZ, schizophrenia; MDD, major depressive disorder; SUD, substance use disorder; OUD, opioid use disorder. Related to **Figure 2**, **Table S13**, **Table S14**.

### Generalized Gene-Set Analysis of GWAS data

Associations of fentanyl self-administration effects in Hb and Amyg with multiple psychiatric and substance use disorders were interrogated using *MAGMA* v1.10 [69]. This method was provided with Single Nucleotide Polymorphisms (SNP) summary statistics from 5 GWAS datasets to compute gene-level associations with the GWAS disorders under the SNP-wise mean Z-statistics method and using the 1000 Genomes European Phase 3 panel [70] as reference. The GWAS datasets were: Panic Disorder (PD) [59], Opioid Use Disorder (OUD) [56], Substance Use Disorder (SUD) [55], and two for Major Depressive Disorder (MDD) [57,58]. Human orthologs of rat Hb and Amyg DEGs between fentanyl vs. saline were obtained using *biomaRt* v2.61.1 [68], and these were subsequently subjected to a competitive positive one-sided gene-set analysis to find associations at the gene-set level according to the gene-level associations (**Figure 4**).

### Software

All the analysis code is available at https://github.com/LieberInstitute/fentanyl_rat_hb_amy [71]. Analyses were performed using R versions 4.3.2 to 4.4.0 [72] with Bioconductor versions 3.17 to 3.19 [73]. Visualizations were made using *ggplot2* v3.4.4 and v3.5.1 [74].

## Results

### Chronic long-access (LgA) fentanyl self-administration results in escalation of intake

A week after intravenous catheter implantation, adult male rats were allowed to self-administer saline (n = 11) or fentanyl (n = 8) in 6 short access sessions (ShA, 2 hours) followed by 16-18 long-access sessions (LgA, 6 hours) (**Figure 1A**). Active lever press resulted in infusion of 0.5 µg of fentanyl dissolved in 50 µl sterile saline, or saline alone (**Figure 1B**). Sixty to ninety minutes following the final LgA phase, brains were collected for gene expression analysis in Hb and Amyg (**Figure 1C**). Thus, the molecular changes we present here reflect the combination of same-day and chronic fentanyl.

As expected, during the LgA phase, fentanyl rats showed higher engagement with the active lever compared to the inactive lever (**Figure S1A-B**), resulting in a greater number of fentanyl infusions and larger infusion slopes compared to saline control rats (**Figure 1D-E**). This pattern was consistent with results from the escalation ratio, in which we normalized the number of infusions obtained per rat across LgA sessions to the number of infusions from the first session (**Figure S1C**). Fentanyl escalation across sessions also remained evident when considering only the first hour of each 6-hour LgA session (**Figure 1F**). On average, rats cumulated a total of 798 µg of fentanyl across LgA sessions (**Figure 1G**). When combining both short- and long-access phases, total intake ranged from 262 to 1,105 µg of fentanyl per rat (**Figure 1G** insert). These results demonstrate a progressive escalation of fentanyl intake with repeated exposure.

We next examined the “futile” lever presses, defined as non-reinforced active lever presses made during the tone and timeout periods (**Figure 1B**). Compared to saline, rats in the fentanyl group consistently made more lever presses during both periods (**Figure S1D-E**). As these lever presses are not reinforced, they may reflect to some extent an increased motivation to obtain the drug with chronic exposure.

### Chronic fentanyl intake is associated with both shared and unique gene expression changes in Hb and Amyg

The Hb and Amyg are key reward circuitry nodes that have been understudied in the context of molecular adaptations following chronic volitional opioid exposure. To identify transcriptomic changes driven by escalating fentanyl self-administration in these regions, we performed bulk RNA-seq in the Hb and Amyg from individual rat brains collected ∼90 minutes after the final LgA session (**Figure 1A**). Following quality control analysis (**Figure S2** to **Figure S7**), we included 16 Hb (8 saline; 8 fentanyl) and 17 Amyg (9 saline; 8 fentanyl) samples in downstream analyses. We analyzed Hb samples separately from Amyg samples to assess correlations between biological, behavioral, and technical variables, and then performed gene expression variance partitioning to build parsimonious linear models for differential gene expression (DGE) analyses (**Figure S8**).

Region-specific analyses between saline and fentanyl treated rats revealed 453 differentially expressed genes (DEGs) in the Hb (271 upregulated, 182 downregulated) and 3,041 DEGs in the Amyg (1,988 upregulated, 1,053 downregulated) at FDR < 0.05 (**Figure 2A**, **Figure S9**, **Table S5**, **Table S6**). In the Hb, top upregulated genes, such as *Cdh10*, *Cttnbp2*, and *Cacna1i*, are mainly involved in synaptic processes and calcium-dependent signaling [75–77] (**Figure 2A**, **Figure S9**). In contrast, in the Amyg, top upregulated genes, including *Mal2*, *Cck*, *Snca*, are linked to vesicle trafficking, mitochondrial organization, and ATP synthesis [78–80] (**Figure 2B**, **Figure S9**). Among the most significant Hb downregulated genes, *Cspg4b* contributes to oligodendrocyte precursor cell structure and myelination [81], *Gstp1* is central to oxidative stress defense and detoxification [82], and *Rnf213* participates in vascular homeostasis [83] (**Figure 2A**). In Amyg, the most significant downregulated genes included *Cep112*, involved in maintenance of centrosome integrity [84], *Abca1*, which mediates cholesterol transport and lipid homeostasis [85], and *Nfatc2*, which regulates calcium-dependent transcription and neuroimmune signaling [86] (**Figure 2B**).

Shared gene expression changes across Hb and Amyg were found (106 shared DEGs), with all but 6 DEGs (*Dact3*, *Plcb4*, *Adarb2*, *Col4a3*, *Abhd17c*, and *Ly6e*) regulated in the same direction (**Table S7**). Notably, *Col4a3* encoding the ⍺3 chain of type IV collagen, which is involved in extracellular matrix (ECM) organization and vascular integrity [87], was oppositely regulated in Hb and Amyg (**Table S7**), suggesting possible region-specific vascular remodeling. Among the top shared DEGs (**Figure 2C**, **Table S7**) were the stress-related receptors *Crhr2* (downregulated) and *Npy1r* (upregulated), whose opposing changes may suggest acute modulation of stress pathways and compensatory, stress-buffering responses following fentanyl intake. Upregulation of the G protein–activated inwardly rectifying potassium (GIRK) channel subunits *Kcnj3* and *Kcnj9*, suggests an increased inhibitory tone and reduced neuronal excitability within the Hb–Amyg network, immediately after fentanyl self-administration. This pattern aligns with evidence that knockout of these subunits alleviates withdrawal symptoms (see [88] for review), suggesting their upregulation reflects a compensatory adaptation to sustained opioid intake.

Next, we performed functional enrichment analyses using Gene Ontology (GO) terms and Kyoto Encyclopedia of Genes and Genomes (KEGG) pathways to identify biological functions linked to both unique and shared Hb and Amyg DEGs (**Figure 2D, Figure S10, Table S8**, **Table S9**). In the Hb, uniquely upregulated genes were primarily associated with synaptic transmission and ionic conductance, including regulators of postsynaptic membrane potential such as *Grik1*, *Scn1a*, and *Gria4*, and voltage-gated cation channels including *Cacna1*, *Kcnj10*, and *Kcnq3* (**Figure 2D**). In contrast, Amyg-specific upregulated genes were enriched for metabolic and vesicular functions, involving mitochondrial and respiratory components such as *Snca*, *Cox6b1*, and *Uqcrq*, together with vesicle-associated genes like *Calm1* and *Arpc3* (**Figure 2D**), consistent with increased energy demand and synaptic output. Shared upregulated genes across both regions converged on synaptic organization and excitability, including *Kcnc2*, *Kctd16*, *Cdh10*, and *Epha4* (**Figure 2D**), reflecting coordinated enhancement of synaptic plasticity within the two structures after fentanyl self-administration. In both regions, fentanyl was also associated with downregulation of extracellular matrix (ECM) genes, such as *Sox9*, *Gsn*, and *Col9a3* (**Figure 2D**), pointing to reduced extracellular remodeling and glial cell support processes (**Table S8**, **Table S9**).

The KEGG pathway enrichment analysis complemented GO results by revealing additional region-specific functional specializations (**Figure S10**). In the Hb, uniquely upregulated genes were enriched for morphine addiction (*Adora1*, *Gabra4*, *Adcy8*), consistent with enhanced opioid-related signaling within this structure. In contrast, the Amyg showed upregulation of genes linked to oxidative phosphorylation (*Atp6v1e1*, *Atp5mc2*, *Atp6v0e2*), Parkinson disease (*Snca*, *Calm1*, *Slc39a10*), and regulation of actin cytoskeleton (*Arpc3*, *Nckap1*, *Cfl1*), indicating increased mitochondrial metabolism and structural plasticity. However, other Amyg-specific downregulated genes within the same pathway (*Vcl*, *C6*, *Wasf2*) suggest a simultaneous suppression of cytoskeletal reorganization, pointing to a complex balance between synaptic remodeling and stabilization following fentanyl exposure. Finally, pathway analyses of shared DEGs revealed opposite regulation of *Col4a3* within ECM–receptor interaction and *Plcb4* across glutamatergic synapse and gastric acid secretion, highlighting region-specific transcriptional divergence in extracellular and excitatory neurotransmission-related signaling between the Hb and Amyg (**Table S8**, **Table S9**).

In addition, we examined transcriptional changes associated with behavior-related metrics among fentanyl rats. We performed DGE for the slope in the number of infusions in the first hour of each LgA session, total overall fentanyl intake across LgA sessions, and fentanyl intake in the last LgA session (**Table S1, Table S2, Table S10**, **Figure S8**). No genes were found significantly associated with these behavioral metrics in either the Hb or Amyg. This is likely due to limited statistical power and higher variability when subsetting to the 8 fentanyl rats.

### Enrichment of genes associated with fentanyl intake in human and rodent Hb and Amyg cell types

Bulk RNA-sequencing is a powerful, high throughput approach to identify gene expression changes in individual animals across experimental conditions. However, it is difficult to identify cell type-specific transcriptomic changes from homogenate tissue profiles, which may be important for understanding the molecular impacts of fentanyl in Hb and Amyg. Therefore, to infer cell types most impacted by fentanyl self-administration in these regions, we performed enrichment analyses leveraging 4 publicly available single-cell and single-nucleus RNA-seq datasets across control rodent and human Amyg and Hb [51–54]. Briefly, the top 100 marker genes for broad and fine-level cell types in each dataset were identified as those with the greatest mean expression in the target cell type compared to any other cell type (**Table S11**, **Table S12**, **Table S13**, **Table S14**). Subsequently, we obtained the rat orthologs of human and mouse cell type markers and assessed their over-representation among fentanyl vs. saline DEGs in rat Hb and Amyg.

First, we assessed enrichment of rat Hb fentanyl-associated DEGs in mouse Hb cell types defined at the broad level by Hashikawa et al. (2020) [51]. We found that upregulated fentanyl DEGs in rat Hb were enriched in a single mouse Hb neuron population (Neuron 8), while downregulated rat fentanyl DEGs were enriched in endothelial cells (**Figure 3A**, **Table S11**). To better understand the identity of Neuron 8, we performed a second enrichment analysis using fine cell type annotations for mouse LHb and MHb subpopulations (**Figure 3B**, **Table S11**) [51]. Upregulated rat Hb fentanyl DEGs were enriched in a specific mouse LHb subpopulation (LHb.6) marked by expression of *Kcnmb4*, *Fam101b*, *Chrm2*, *Sv2c*, and *Gpr151*. We also observed enrichment of downregulated fentanyl DEGs in mouse LHb.2, marked by expression of *Arpp21*, *Cacna2d1* and *Slc6a1*, at this fine cell type resolution. Next, to evaluate whether rat fentanyl DEGs might be enriched in orthologous Hb subpopulations identified in the human brain, we ran an enrichment test across LHb and MHb neuronal populations and glial cell types from our previously published human Hb snRNA-seq dataset [52] (**Figure 3C**, **Table S12**). We found that upregulated and downregulated rat fentanyl DEGs registered to human glial cell types, including astrocytes, oligodendrocytes, and endothelial cells. We did not see fentanyl DEG enrichment in any human Hb neuronal subpopulations, likely due to limitations in power as a result of the small number of LHb and MHb neurons captured in this human dataset. Interestingly, human Hb LHb.2 and LHb.7 subpopulations show high expression of *KCNMB4, GPR151, and CHRM2* **(Figure S11A-C**), suggesting these cell types may be conserved with mouse LHb.6, which showed an enrichment of upregulated fentanyl DEGs (**Figure 3B**, **Table S11**). Human LHb.2 and LHb.7 also highly express *OPRM1* compared to other human LHb subpopulations (**Figure S11D**), suggesting these may be fentanyl-sensitive LHb populations conserved across species.

Next, we performed similar enrichment analyses of rat Amyg fentanyl-associated DEGs in previously identified Amyg cell types from rat [54] and human [53] snRNA-seq datasets. At the broad cell type level when comparing within species, we found that upregulated rat fentanyl DEGs were enriched in rat endothelial cells and excitatory neurons, while downregulated fentanyl DEGs were enriched in oligodendrocytes (**Figure 4A**, **Table S13**). At the fine cell type level, upregulated fentanyl DEGs were still most strongly enriched in excitatory neurons, while downregulated DEGs remained enriched in oligodendrocytes and oligodendrocyte precursor cells (OPCs). However, at the finer level, we also observed enrichment of upregulated fentanyl DEGs in astrocytes and several inhibitory neuron populations, including *Cck*+/*Vip*+, *Nos1*+, and *Pvalb*+ (**Figure 4B**, **Table S13**). Notably, *Cck* is one of the top 5 upregulated genes following chronic fentanyl intake in Amyg (**Figure 2**). Enrichment analyses in human Amyg cell types largely agreed with findings in rodents. At the fine cell type level, upregulated rat fentanyl DEGs were enriched in Endo_NOSTRIN populations and several different excitatory subpopulations in the basolateral amygdala (BLA) expressing *HGF*, *SATB2*, and *LAMP5* (**Figure 4C**, **Table S14**). Similar to the rodent analysis, upregulated rat fentanyl DEGs were also enriched in several human inhibitory cell types, such as *SST*+ and *VIP*+ subpopulations, and *TSHZ1*+ intercalated neurons. Of particular interest, *TSHZ1*+ intercalated neurons receive input from VTA dopamine neurons and highly express dopamine receptor 1 (*DRD1*) and *OPRM1* [89,90]. Downregulated rat fentanyl DEGs were enriched in several human glial cell types, including specific populations of oligodendrocytes, OPCs, astrocytes, and microglia (**Figure 4D**, **Table S14**).

Finally, we asked whether fentanyl DEGs in Hb and Amyg overlapped with genes associated with risk for various neuropsychiatric disorders, including substance use disorder (SUD) and opioid use disorder (OUD), as identified by genome wide association studies (GWAS) [55–60]. For Amyg, we found that upregulated fentanyl DEGs were enriched in schizophrenia (SCZ)-associated risk genes, while downregulated fentanyl DEGs were enriched in SUD-associated risk genes. No enrichment was observed for fentanyl DEGs in Hb (**Figure 4E**). Taken together, we identify the enrichment of fentanyl-associated genes in rodent and human Hb and Amyg cell types and demonstrate overlap of these genes with risk genes for SUD in Amyg.

## Discussion

While several studies have examined transcriptomic changes associated with opioid exposure across brain reward structures (e.g. NAc, VTA, dorsal striatum, prefrontal cortex), fentanyl-induced transcriptional programs in the Hb and Amyg have not yet been investigated. Here, we established a chronic long-access fentanyl self-administration paradigm in which rats show escalating fentanyl intake. We performed bulk RNA-seq on individual rats following the last self-administration session and identified both unique and shared DEGs in the Hb and Amyg, many of which were implicated in distinct functional pathways. We integrated fentanyl-associated DEGs with sc/snRNA-seq data from rodent and human Hb and Amyg and identified neuronal and glial cell types enriched in fentanyl DEGs. Finally, we assessed the overlap of fentanyl DEGs with genes associated with risk for neuropsychiatric disorders, including SUD, and found enrichment of SUD risk genes among downregulated fentanyl DEGs in Amyg.

Sixty to ninety minutes after the last session of chronic long-access fentanyl self-administration, we observed both distinct and convergent transcriptional adaptations in the Hb and Amyg. In the Hb, we found upregulated genes related to synaptic transmission and ionic conductance, suggesting heightened excitability, whereas downregulated Hb DEGs involved myelination, vascular regulation, and oxidative stress defense indicating weakened structural and metabolic support. The Amyg, by contrast, showed enrichment for mitochondrial activity, ATP synthesis, and vesicular pathways, consistent with increased energy demand and neurotransmitter release, alongside reduced expression of genes governing cytoskeletal organization, lipid metabolism, and neuroimmune balance. Despite some regional differences, both regions exhibited coordinated upregulation of genes promoting synaptic organization and excitability, accompanied by suppression of ECM and glial-related genes. These findings align with postmortem human studies of individuals diagnosed with OUD, which reveal disrupted synaptic plasticity, ECM and neuroimmune-related functions in the PFC and NAc [45]. Additionally, neuroinflammatory and metabolic alterations across neuronal and glial populations were also observed in the dorsal striatum of individuals who died from opioid overdose [46]. Collectively, these findings point to maladaptive synaptic plasticity as well as altered metabolic and glial homeostasis following chronic opioid volitional intake.

The Hb and Amyg fentanyl-induced transcriptional changes we observed in this study closely align with other recent fentanyl transcriptomic studies. Notably, Wood et al. [91] examined DEGs in the rat CeA following fentanyl exposure. Among the 2,735 genes common to their dataset and our Amyg dataset, only seven (*Tuba4a*, *Atrnl1*, *Ift140*, *Arl8b*, *Ndufa10L1*, *Ndrg4*, *Bhlhe22*) were significantly altered in both. These genes are implicated in functions such as cytoskeletal organization, mitochondrial metabolism, and signal transduction. Limited DEG overlap may stem from differences in sample size, but also likely differences in experimental design: Wood et al. investigated perinatal fentanyl exposure followed by adult self-administration (3 h/day for 12 days) with tissue collected 24 h after the final session, whereas our study examined adult-onset, long-access self-administration (2 h/day for 6 days + 6 h/day for 16–18 days) with tissue collected 90 min post-session. Uchegbu and coworkers [92], found that prenatal fentanyl exposure in mice led to upregulation of myelin- and glia-related genes in the Amyg whereas these pathways were downregulated in our dataset, suggesting opposing transcriptional adaptations between developmental and adult exposure. Complementing these transcriptional studies, an *ex vivo* calcium imaging experiment in rat showed that long-access fentanyl self-administration promotes a reduction in central Amyg activity after a 30 days abstinence period and a shift from acute tolerance to chronic hypersensitivity to fentanyl exposure [93]. These neuronal adaptations likely reflect underlying transcriptional remodeling that disrupts Amyg excitability over time and may drive fentanyl withdrawal symptoms and relapse.

Our dataset also partially overlaps with transcriptomic changes in the VTA after chronic fentanyl self-administration [43]. Our Amyg dataset showed a large and concordant overlap with the mouse VTA (285 shared DEGs), with most shared genes downregulated in both regions and involved in metabolic, mitochondrial, and myelin-related pathways, together with a coordinated upregulation of neuronal excitability and synaptic plasticity genes. While Amyg showed broad overlap with the VTA, the Hb exhibited a distinct transcriptional profile. Among the 58 genes significantly altered in both Hb and VTA, a subset showed concordance across regions, with upregulated DEGs showing enrichment for glutamatergic and GABAergic synaptic signaling and synapse organization, and downregulated DEGs enriched for lipid metabolism and ECM, which reflects shared neuroadaptation processes. However, nearly half of the shared DEGs (24 genes) displayed opposite regulation across the two regions, being upregulated in Hb but downregulated in VTA. These included genes involved in synaptic organization, neuronal signaling and cytoskeletal regulation (*Cttnbp2*, *Kcnq3*, *Kcnj10*,*Il1rap*, *Arhgap21*, *Rock1*, *Arap2*), metabolic and mitochondrial regulation (*Pnpla7*, *Slc25a27*, *Hif3a*, *Abcc5*), and protein modification or stress responses (*Usp31*, *Hltf*, *Htra1*). Given species differences and the distinct post-exposure time points across studies (24 h vs 90 min), these comparisons should be interpreted cautiously. Yet, this opposite transcriptional profile may suggest a region-specific action of fentanyl, with enhanced structural plasticity and metabolic demand in the Hb, contrasting with a dampening in the VTA. These molecular results are consistent with photometry recordings of the LHb during oral fentanyl self-administration [94], where progressive increases in drug-evoked LHb activity across three weeks of training suggest plasticity accompanying the transition from positive to negative reinforcement.

A study by Olusakin et al. [95] reported transcriptional adaptations across the mesolimbic pathway (VTA→NAc) following perinatal and juvenile fentanyl exposure in mice. Both VTA and NAc showed strong enrichment for genes linked to mitochondrial respiration, ECM remodeling, and synaptic and structural remodeling, functions that overlap with fentanyl-induced genes identified in our Hb and Amyg datasets. In the VTA, genes such as *Lrrtm2*, *Dynlt3*, and *Ndufaf4* were upregulated, reflecting enhanced synaptic and mitochondrial activity, whereas ECM-related genes (*Col9a3*, *Dact3*) and oxidative stress regulators (*Selenom*) were downregulated, indicating selective suppression of matrix and antioxidant processes. However, we did not observe significant overlap between our DEGs and those altered in the NAc in Olusakin and colleagues’ dataset. In addition, although no common DEGs were identified between our Hb and Amyg datasets and the mouse NAc profiles reported by Fox et al. [96], their data revealed mitochondrial, extracellular matrix, and synaptic GO term enrichments that match those dysregulated in our study. The absence of direct gene-level overlap likely reflects differences in species, brain regions, or experimental paradigms, with short oral fentanyl exposure and a long withdrawal period in their study, whereas our approach involved chronic fentanyl i.v self-administration and immediate DEGs assessment. Together, these findings support convergent, yet region-specific transcriptional adaptations in synaptic, metabolic, and structural processes across reward and aversion circuits following fentanyl exposure. These results are further confirmed by a recent study [44] where escalating fentanyl intake in long-access sessions was related to gene expression changes in calcium and potassium channels as well as synaptic signaling in the mPFC.

This transcriptional pattern extends beyond fentanyl self-administration and appears to be conserved across opioids and opiates. Indeed, morphine self-administration similarly enhanced synaptic, vesicular, and mitochondrial functions in the NAc while reducing extracellular matrix, vascular, and glial-related genes [40]. Likewise, heroin exposure produced comparable changes across several mesocorticolimbic regions, ventral hippocampus and BLA [9], reflecting a general shift toward heightened neuronal excitability and metabolic demand under diminished cellular support induced by mu receptor agonists.

To better understand which cell types are impacted by chronic fentanyl self-administration, we integrated our fentanyl DEGs with sn/scRNAseq data from rodent and human Hb and Amyg. This allowed us to overcome limitations in cell type heterogeneity in our bulk RNA-seq data, while leveraging the power of this approach to profile transcripts across different cell compartments, including the nucleus, cytoplasm, neuronal processes, and synapses. Consistent with previous sc/snRNA-seq studies that identified opioid-related transcriptional changes in glial populations in the rodent NAc and Amyg [30,34,40], we found enrichment of our fentanyl DEGs in oligodendrocytes, astrocytes and endothelial cells in both Hb and Amyg. These findings were conserved across species, and highlight that fentanyl impacts cell types critical for blood brain barrier (BBB) integrity and myelination. Notably, the BBB is an area of therapeutic interest as it regulates fentanyl access to the central nervous system [97], and opioids alter BBB homeostasis, structure, and integrity [98–100]. Further supporting a role for fentanyl in oligodendrocyte dysfunction, myelin pathology has been implicated in OUD [101,102] and chronic methadone treatment of primary rat glia cultures increases oligodendrocyte apoptosis and reduces myelinating capacity [103]. Future functional studies should investigate how fentanyl intake dysregulates the neurovascular unit.

Beyond glial cells, we also observed enrichment of upregulated and downregulated rat Hb fentanyl DEGs in specific mouse LHb cell types identified by Hashikawa et al. [51]. In particular, upregulated DEGs were enriched in mouse LHb.6, which is marked by expression of *Kcnmb4*, *Gpr151*, and *Chrm2*. These genes are also highly expressed in two *OPRM1*-expressing human Hb populations, LHb.2 and LHb.7 [52], suggesting possible conservation of fentanyl-sensitive mouse LHb.6 with human *OPRM1*-expressing LHb subpopulations. Notably, foot shock induces activity-dependent gene expression in LHb.6 [51], implicating this population in the processing of painful noxious stimuli and providing further support for the functional relevance of this LHb cell type in opioid signaling. Unfortunately, direct enrichment analyses of rat Hb fentanyl DEGs in our human Hb cell types did not yield any significant results, likely due to a lack of power resulting from the limited number of human Hb cells in our dataset.

In the Amyg, upregulated rat fentanyl DEGs were enriched in several excitatory and inhibitory populations, whereas downregulated DEGs were exclusively enriched in glial cell types, especially oligodendrocytes, which is consistent with findings in rodent and human Hb. We observed enrichment of upregulated rat fentanyl DEGs in multiple human fine level excitatory Amyg cell types, including *SATB2*+, *HGF*+, and *LAMP5*+ subclusters, which likely represent BLA excitatory populations. This is consistent with previous work showing that BLA glutamatergic principal neurons have increased intrinsic excitability and excitatory/inhibitory ratio in mice exposed to oral fentanyl consumption and withdrawal [24]. Future work should investigate the relationship between molecular and physiological changes in BLA pyramidal populations following chronic fentanyl intake and withdrawal. In terms of GABAergic neurons, vasoactive intestinal peptide (*VIP*)+ interneurons and *TSHZ1*+ intercalated neurons showed enrichment in upregulated genes in both rodents and humans. In the hippocampus, cortex, and striatum, there is mounting evidence for interactions between *VIP*+ interneurons and the opioid system, with *VIP*+ neurons showing *Oprm1* expression across many brain regions [104,105]. *TSHZ1*+ intercalated neurons are highly conserved across species [90], densely express *Oprm1,* and act to inhibit BLA and CeA circuitry under basal conditions [106,107]. However, *Oprm1* activation leads to inhibition of intercalated neurons, thus disinhibiting BLA and CeA reward circuitry underlying relapse of drug seeking behavior [108]. Together, these findings point towards a conserved opioid-driven disruption of glial support and inhibitory microcircuits that shifts amygdala function toward the disinhibition of neural circuits underlying relapse.

We acknowledge the present study has several limitations that set the stage for future investigations. First, we only perform molecular profiling at one timepoint ∼90 minutes following the last self-administration session. We chose this time point to capture the transcriptomic changes following excessive chronic fentanyl use while avoiding transcriptomic changes induced by acute withdrawal. Hence, it will be important to distinguish the changes we observed from those following a single acute administration of fentanyl, although we expect distinct changes given prior comparisons of gene regulation after acute and chronic drug [9,109]. Additionally, it will be important to look at fentanyl-associated changes at other timepoints across different phases of opioid self-administration, including across acquisition, withdrawal and relapse. Given that the Hb is implicated in opioid withdrawal [110–112], future studies should assess transcriptomic changes following an abstinence period. Such changes could underlie alterations in risky decision making observed during fentanyl abstinence [113], and potentially contribute to relapse vulnerability. Second, while bulk RNA-seq is high throughput and cost-effective, higher resolution technologies such as snRNA-seq will be needed to validate inferred cell type-specific changes associated with chronic volitional fentanyl intake. As discussed above, snRNA-seq studies in the rodent NAc, VTA, and Amyg following opioid administration [34,40,43] have yielded valuable insights into neuronal and glial populations impacted by fentanyl and other opioids. Third, we acknowledge limitations in power that prevented the identification of DEGs associated with behavioral measures in fentanyl-administering rats, including total fentanyl intake. Future experiments with larger numbers of rats should leverage more complex behavioral analyses using tools, such as machine learning-based behavioral tracking [114], for integration with transcriptomic data. Furthermore, given previously reported sex differences in humans and rodents in the context of opioids [42,115–117], future studies should also include female rats as we were only able to assess males in the present study. Finally, it will also be important to consider different control groups, such as a natural reward group, to identify gene expression changes unique to an addictive drug like fentanyl from those induced by a non-addictive reward like sucrose.

In summary, we report transcriptional programs associated with fentanyl intake in the rat Hb and Amyg and link fentanyl-associated gene expression changes to specific cell types in the rodent and human brain. We provide evidence that genes altered by fentanyl in the rat Amyg overlap with genes implicated in genetic risk for neuropsychiatric disorders, including substance use disorders. Taken together, these findings highlight cell types and molecular pathways for further functional follow that could be targeted for potential therapeutic development.

## Acknowledgements

We gratefully acknowledge use of the facilities at the Joint High Performance Computing Exchange (JHPCE) in the Department of Biostatistics, Johns Hopkins Bloomberg School of Public Health that have contributed to the research results reported within this paper. We thank Keri Martinowich for helpful comments on the manuscript.

## Conflict of Interest

The authors declare no conflicts of interest.

## Supplementary Methods

### Library construction and bulk RNA sequencing (RNA-seq)

Total RNA was extracted from habenula and amygdala tissue samples using the Qiagen RNeasy Micro Kit (Cat. No.: 74004). Paired-end strand-specific sequencing libraries were prepared for 33 samples (16 habenula, 17 amygdala) from 100 ng total RNA input for amygdala samples, and 10 ng total RNA input from low-yield habenula samples. For amygdala samples, TruSeq Stranded Total RNA Library Preparation kit with Ribo-Zero H/M/R Gold ribosomal RNA depletion was used (min. 0.5 ug of total RNA needed per sample). For low-yield habenula samples, Illumina Stranded Total RNA Prep Library Preparation kit with Ribo-Zero Plus was used (∼1 ng to 1000 ng of total RNA needed per sample). For quality control, synthetic External RNA Controls Consortium (ERCC) RNA Mix 1 (Thermo Fisher Scientific) was spiked into each sample. Libraries were sequenced on an Illumina NovaSeq 6000 S4 (152 bp PE) producing ∼80 million (median 102,396,054, mean 98,533,714) 152 bp paired-end reads per sample.

### RNA-seq data processing

#### Gene expression quantification

*SPEAQeasy* pipeline version 87ba0b4 [63], a *Nextflow* v20.01.0 [118] workflow for *HISAT2* v2.2.1 [119], was used with default settings to assess the quality of the sequencing reads and quantify the gene expression in the samples using the rat genome assembly mRatBN7.2 from Ensembl release 109 [120,121]. A *RangedSummarizedExperiment* R object [73] with gene counts for 30,452 genes across 33 samples was built by *SPEAQeasy*; this object included sample quality metrics that were used for exploratory analyses (**Supplementary Methods: Exploratory Data Analysis**).

#### Filtering of lowly-expressed genes

Lowly-expressed genes were filtered using `filterByExpr()` from *edgeR* v3.43.7 [122] in which only genes with at least 15 total reads across all samples and with 10 or more counts in at least *n* samples are retained, where *n* is defined as 70% the size of the smallest sample group. After this step, 16,708 genes (54.86%) were retained for downstream analyses.

#### Count normalization

Raw expression counts of the genes in the 33 samples were normalized by trimmed mean of M-values (TMM) [64] using `calcNormFactors()` from *edgeR* v3.43.7 [122] to compute normalization factors for library size rescaling. *edgeR* `cpm()` [122] was subsequently used to obtain counts per million (CPM) in a logarithmic scale: approximately log_2_(CPM+0.5).

### Exploratory Data Analysis (EDA)

#### Sample Quality Control Analysis (QCA)

Sample-level gene-based quality control (QC) metrics computed by *SPEAQeasy* [63] on raw counts before gene filtering and count normalization steps (**Table S3**, **Table S4**), were compared across the different brain regions, substances, preparation batches, and rat administration sessions. Hb samples presented lower yields of RNA compared to Amyg samples, which in turn decreased their library sizes, number of detected genes, and read mapping rates (**Figure S2**). The third RNA extraction batch was performed on additional samples and resulted in more comparable yields with the second higher-yield Amyg batch. Hb and Amyg samples from the third RNA extraction batch presented good RNA amounts and library sizes, and higher mapping rates than their counterparts in the first and second batches, respectively (**Figure S3**). No association was observed between sample quality metrics and total number of fentanyl self-administration sessions (**Figure S4**). Hb and Amyg samples were analyzed separately in subsequent steps.

Low-quality samples were defined through the identification of outlier QC metrics with `isOutlier()` from *scater* v1.30.1 [123], which takes as outliers those values that are 3 median-absolute-deviations (MAD) away from the median. For Hb, one of the 16 samples was detected as an outlier for the number of detected genes (**Figure S5A**) and for Amyg, 3 of the 17 samples were detected as outliers for mitochondrial mapping rate, concordant mapping rate, or number of detected genes (**Figure S5B**); two of the three Amyg outliers were from the third RNA extraction batch. However, these four samples had no other outlier QC metrics (**Figure S5**) and were not removed but examined further in principal component (PC) plots (**Supplementary Methods: Sample-level gene expression variation and manual QC inspection**). Sample QC metrics are described in **Table S3**.

#### Sample-level gene expression variation and manual QC inspection

Sources of gene expression variation between samples were explored with Principal Component Analysis (PCA) on log-normalized counts of expressed genes. RNA extraction batch appeared as a major driver of gene expression variability in both Hb and Amyg (**Figure S6**), and substance had a greater contribution among Amyg samples (**Figure S6B**). The four previous outlier samples in QC metrics (**Figure S5**) were not outliers on PC plots, although other samples were detected as outliers by PCA (**Figure S7**). Both QC metrics and PCA outlier samples were subjected to manual inspection of all their QC metrics, finding only attenuated differences in the quality of these samples compared to the non-outlier ones, as well as high-quality metrics for these samples relative to the global metrics estimates (**Figure S7**). All Hb and Amyg samples were retained for posterior analyses.

#### Gene-level expression variation and covariate selection for DGE

To explore the contributions of sample-level variables on gene expression variation and guide variable selection to model gene expression for DGE analysis, we first computed the percentage of variance of gene expression explained by each covariate individually with `getVarianceExplained()` from *scater* v1.30.1 [123]. We implemented this analysis taking all Hb and Amyg samples separately (**Figure S8**) and then subsetting to fentanyl-administered samples from each brain region (**Figure S8**), as additional DGE analyses were performed on fentanyl-administered samples only (see further below and **Supplementary Methods: Differential Gene Expression analysis**).

Second, for the same sample groups, we performed pairwise Canonical Correlation Analysis (CCA) with `canCorPairs()` of *variancePartition* v1.32.5 [124] to identify pairs of correlated variables (**Figure S8**). To remove redundant and minority contributing variables and to avoid unmasking true drivers of variation, models for DGE between fentanyl vs. saline administration and for rat behavior in Hb and Amyg were defined by discarding:

1. variables highly correlated with “substance” (fentanyl vs. saline; **Figure S8**) or “behavioral covariates” (i.e. total fentanyl intake, last session fentanyl intake, or slope for the rats’ fentanyl intake in the first hour of each session; **Figure S8**);
2. variables highly correlated with the RNA extraction batch previously shown to affect samples’ quality metrics (**Figure S3**) and explain high percentages of expression variation in several genes (**Figure S8**);
3. variables highly correlated with the total number of fentanyl sessions and this variable itself, as it didn’t impact on sample QC metrics (**Figure S4**) and had minor contributions on gene expression differences (**Figure S8**), and
4. for other pairs of correlated variables, we only kept the one with the highest percentages of gene expression variance explained obtained with `getVarianceExplained()` (**Figure S8**).

Then, the fraction of variation in the expression of each gene attributable to each included variable was assessed with a variance partition analysis using `fitExtractVarPartModel()` from *variancePartition* [124], jointly accounting for the contributions of the rest of selected variables to confirm their impacts on gene expression and suitability for DGE (**Figure S8**). Sample variables are defined in **Table S3**.

#### Differential Gene Expression (DGE) analysis

We assessed DGE for substance and behavior under the empirical Bayes framework of *limma-voom* v3.58.1

1. [65] pipeline, fitting a linear model to the expression of each gene including as covariates the sample variables that were not correlated between them and that explained high percentages of global gene expression variance (**Supplementary Methods: Exploratory Data Analysis**). The gene-wise *p*-values of the resulting moderated *t*-statistics were adjusted for multiple testing using the Benjamini and Hochberg’s (BH) procedure that controls the false discovery rate (FDR) [125]. Genes with FDR adjusted *p*-values below 0.05 were considered as DEGs.

The following were the specific DGE analyses performed and the covariates included to model gene expression in each.

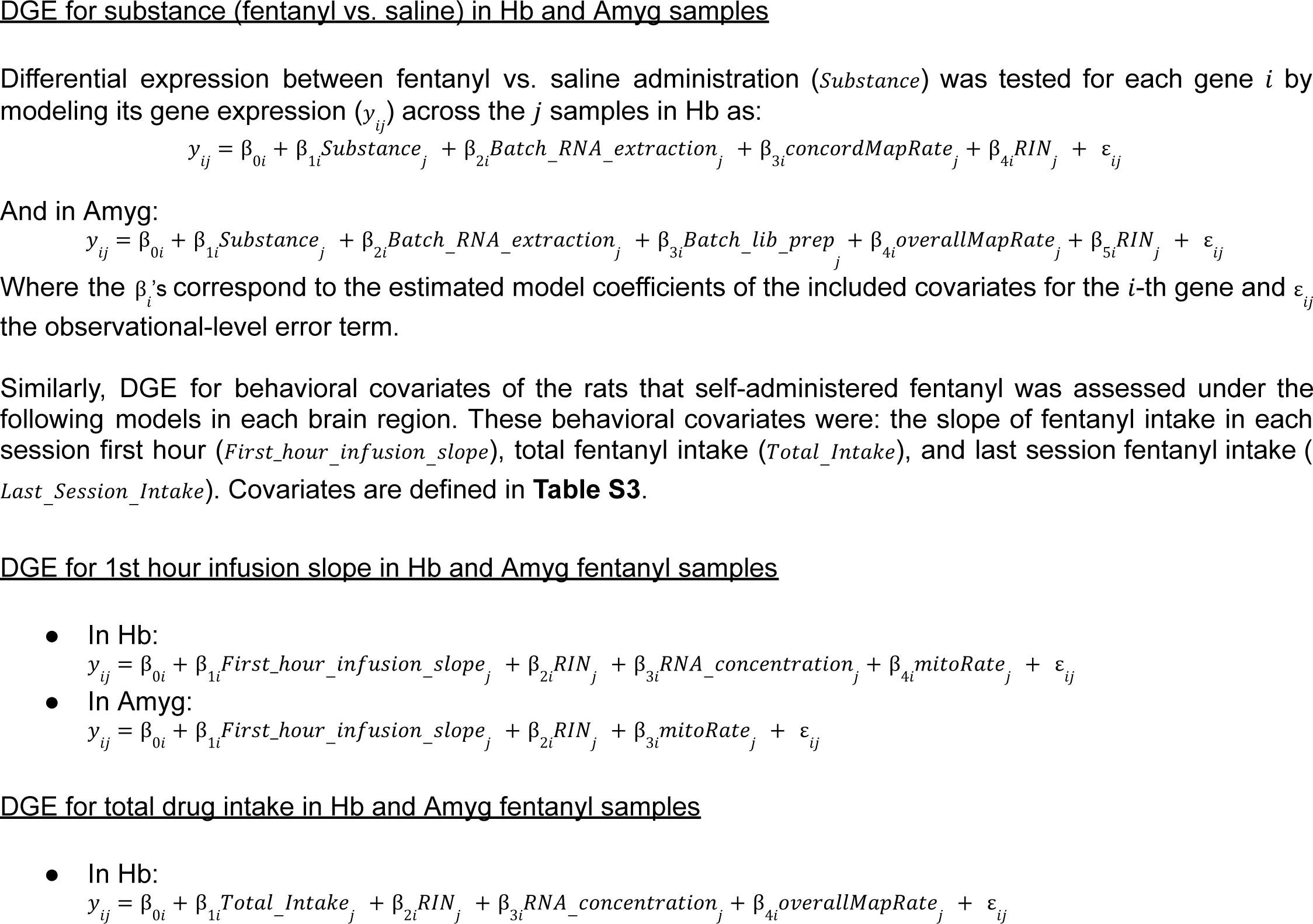
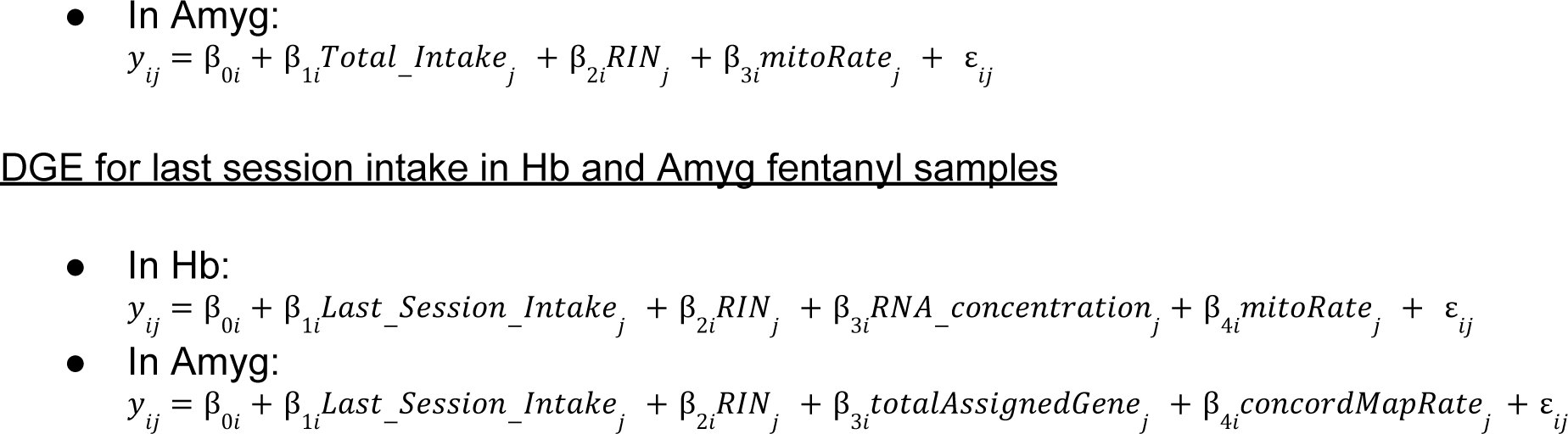

#### Functional enrichment analysis

Gene sets annotated in Gene Ontology (GO) [126] terms and Kyoto Encyclopedia of Genes and Genomes (KEGG) [127] pathways that were significantly overrepresented among our Hb and Amyg DEGs for substance were found with hypergeometric tests implemented in *clusterProfiler* v4.10.0 [66] using `compareCluster()`. The complete set of expressed genes that were assessed for DGE and with available Entrez gene IDs (n=14,066 genes) was taken as the background gene set. The obtained gene set *p*-values were FDR-adjusted [125].

#### Cell type enrichment analysis

Marker genes for 1) main cell types and and inhibitory neuronal subtypes in control rat Amyg [54], 2) cell types at fine and broad resolutions in the human Hb-enriched epithalamus [52] and human Amyg [53] of neurotypical control donors donors, and 3) all and the Hb neuronal cell subpopulations in the Hb complex of control mice [51] were obtained using normalized and filtered sn/scRNA-seq data. Markers were found implementing the *MeanRatio* method of *DeconvoBuddies* v0.99.0 [67]. Cell types with less than 10 cells were discarded from marker finding analysis.

Briefly, *MeanRatio* defines as cell type markers those genes with the greatest mean expression in the target cell type compared to any other cell type, computing for each gene the ratio between the mean expression in the target cell type, and the highest mean expression among the non-target cell types (i.e. the *MeanRatio*) [67]. The top 100 or 50 (human Hb [52]) marker genes with *MeanRatios* >1 were used (**Table S11**, **Table S12**, **Table S13**, **Table S14**).

Then, rat orthologs of human and mouse cell type marker genes were obtained using *biomaRt* v2.56.1 [68] under the GRCh38 and GRCm39 genome versions for human and mouse, respectively. The sets of cell type-specific markers in rat were assessed for their enrichment among all, up-, and down-regulated fentanyl vs. saline DEGs in rat Hb and Amyg based on the one-sided Fisher’s exact test. Expressed genes assessed for DGE were considered the background gene set (n=16,708 genes).

#### Generalized Gene-Set Analysis of GWAS data

*MAGMA* v1.10 [69] was run to assess the joint association of genes in each set of all, up-, and down-regulated Hb and Amyg DEGs with multiple SUDs and psychiatric disorders. Briefly, *MAGMA* was provided as input the summary statistics of genome-wide human Single Nucleotide Polymorphisms (SNPs) from six GWASes: schizophrenia (SCZ) [60], Panic Disorder (PD) [59], Opioid Use Disorder (OUD) [56], Substance Use Disorder (SUD) [55], and Major Depressive Disorder (MDD) [57,58] .

For each GWAS, autosomal SNPs were first mapped onto human genes based on the same human genome reference build used in each study (either GRCh37 or GRCh38; hg19 or hg38). For the gene-level analysis SNP *p*-values were used to compute gene-level *p*-values for their association with the phenotype through the SNP-wise mean Z-statistics method. Given that most of the ancestry composition of all examined GWASes was of European Ancestry, the 1000 Genomes European Phase 3 panel [70] was used as the reference dataset to account for linkage disequilibrium between SNPs.

For gene-set analysis, the human orthologs of rat Hb and Amyg DEGs from each set (all, up-, and down-regulated) were first retrieved from Ensembl release 112 (mRatBN7.2) [128] using *biomaRt* v2.61.1 [68]. Then a competitive positive one-sided gene-set analysis was implemented on such sets of human orthologs to assess their association with the GWAS phenotype based on gene-level associations.

## Author Contributions

Conceptualization: DGP, EAY, ECP, PHJ, LCT, KRM Software: DGP, LCT

Formal Analysis: RM, DGP, EAY, ECP, NJE Investigation: EAY, ECP, PHJ, KRM Resources: PHJ, LCT, KRM

Data Curation: RM, DGP, EAY, MST

Writing-original draft: RM, DGP, EAY, PHJ, LCT, KRM

Writing-review and editing: RM, DGP, EAY, ECP, PHJ, LCT, KRM Visualization: RM, DGP, EAY

Supervision: PHJ, LCT, KRM

Project administration: PHJ, LCT, KRM Funding Acquisition: PHJ, LCT, KRM

PHJ, LCT, and KRM had full access to all the data in the study and take responsibility for the integrity of the data and the accuracy of the data analysis.

## Funding/Support

This work was supported with funding from the Lieber Institute for Brain Development and National Institutes of Health (NIH US) grants T32MH015330 (Yalcinbas), R21DA060407 (Maynard), and R01DA035943 (Janak).

## Data Sharing Statement

The source FASTQ files are publicly available from the NCBI Sequence Read Archive BioProject PRJNA1179901. All analysis code is available at https://github.com/LieberInstitute/fentanyl_rat_hb_amy.

## Supplementary Figures

**Figure S1:**
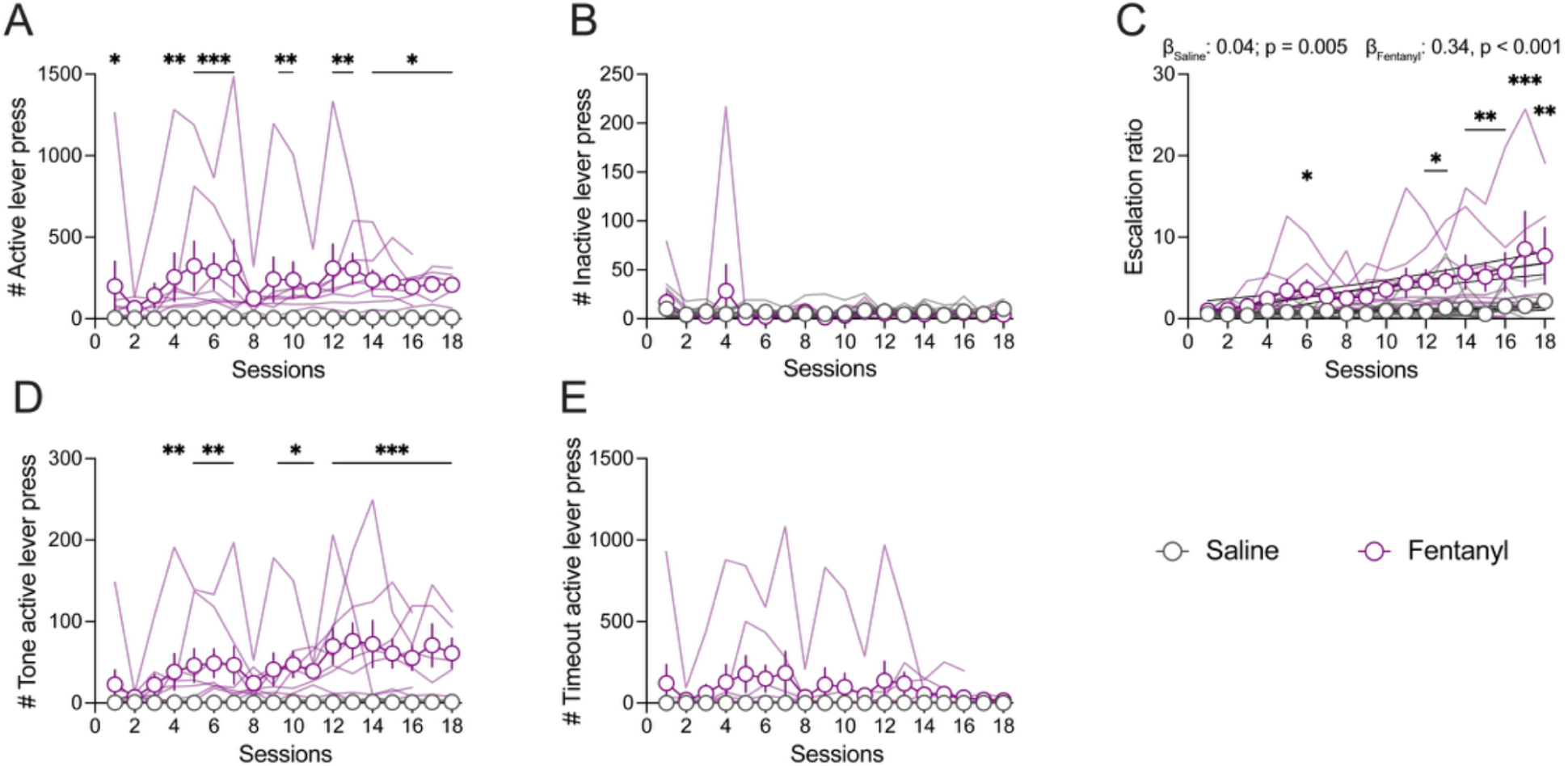
Additional behavior metrics from LgA sessions. Mean number of total (**A**) active (two-way RM ANOVA, substance effect: F_1,17_ = 11.55, p = 0.003; session effect: F_17,273_ = 1.44, p = 0.115; substance x session interaction: F_17,273_ = 1.4, p = 0.133; Student Newman-Keuls post hoc pairwise comparison; ANCOVA substance slope difference: F_1,322_ = 32.07, p < 0.001) and (**B**) inactive lever presses per LgA session for saline and fentanyl rats (two-way RM ANOVA, substance effect: F_1,17_ < 1, p = 0.913; session effect: F_17,273_ = 1.27, p = 0.214; substance x session interaction: F_17,273_ = 1.2, p = 0.265; ANCOVA substance slope difference: F_1,322_ = 32.07, p < 0.001). (**C**) Escalation ratio, an alternative metric to quantify infusion escalation across sessions. This ratio is calculated by normalizing each rat’s LgA infusion counts relative to their infusion count on the first LgA session (two-way RM ANOVA, substance effect: F_1,17_ = 8.49, p = 0.01; session effect: F_17,273_ = 4.66, p < 0.001; substance x session interaction: F_17,273_ = 2.8, p < 0.001; ANCOVA substance slope difference: F_1,322_ = 22.88, p < 0.001). (**D**) Number of active lever presses performed during the 2.8 second infusion and tone presentation period (two-way RM ANOVA, substance effect: F_1,17_ = 19.68, p < 0.001; session effect: F_17,273_ = 2.9, p < 0.001; substance x session interaction: F_17,273_ = 2.8, p < 0.001). (**E**) Number of active lever presses performed during the 20 second timeout period (two-way RM ANOVA, substance effect: F_1,17_ = 3.25, p = 0.089; session effect: F_17,273_ = 1.4, p = 0.132; substance x session interaction: F_17,273_ = 1.4, p = 0.13). Data shown as mean across rats ± SEM, superimposed with individual rat data points. Black lines represent linear regression with 95% confidence intervals. Saline: n = 11 rats; Fentanyl: n = 8 rats. *p < 0.05; **p < 0.01; ***p < 0.001.

**Figure S2:**
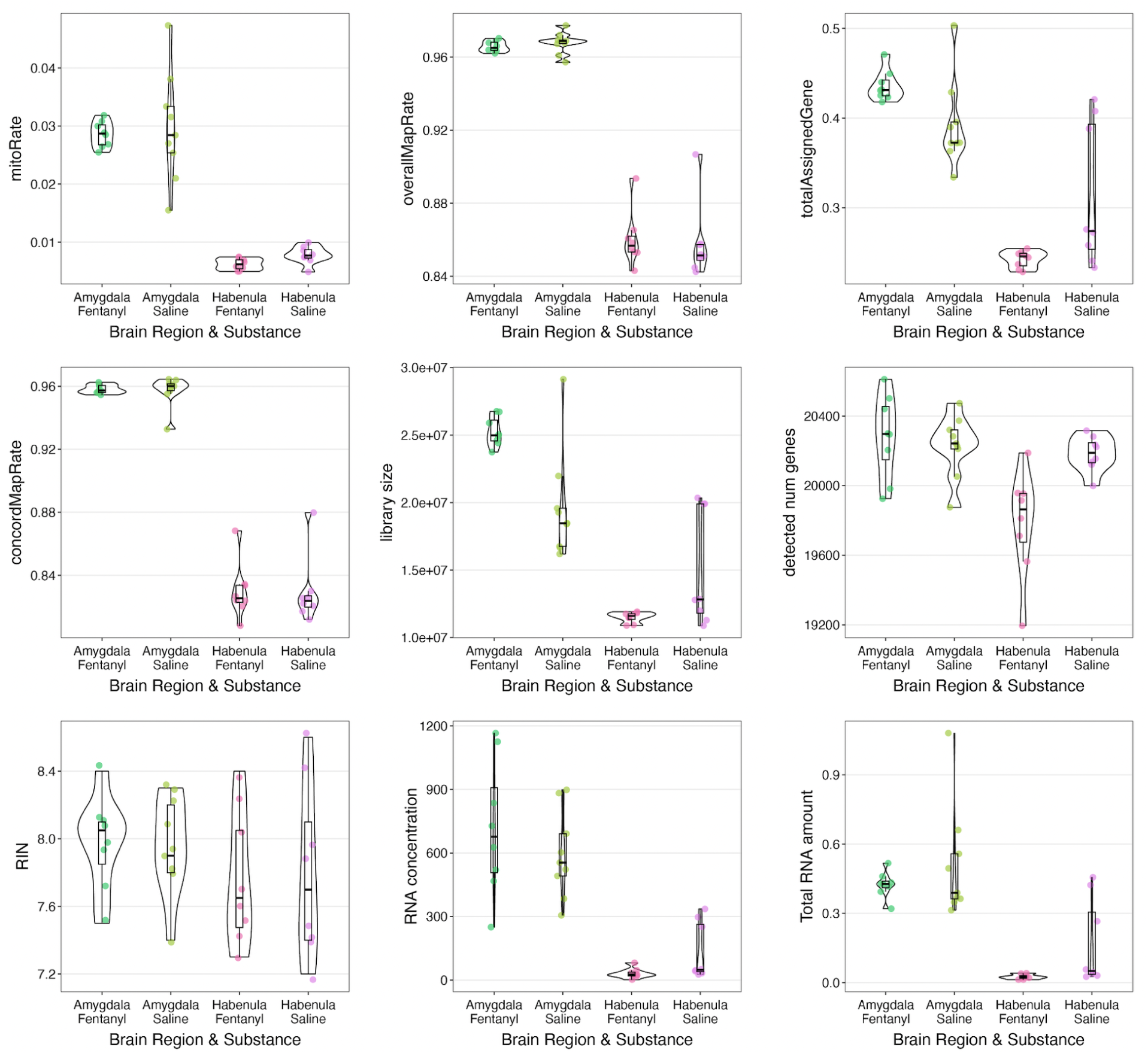
Quality control metrics for Hb and Amyg samples. Comparison of the QC metrics examined in this study for habenula and amygdala fentanyl and saline samples. Note that different Illumina library preparation kits were used for each brain region, thus confounding brain region and kit differences, which motivated independent analyses for each brain region. See **Table S3** for the description of these QC metrics.

**Figure S3:**
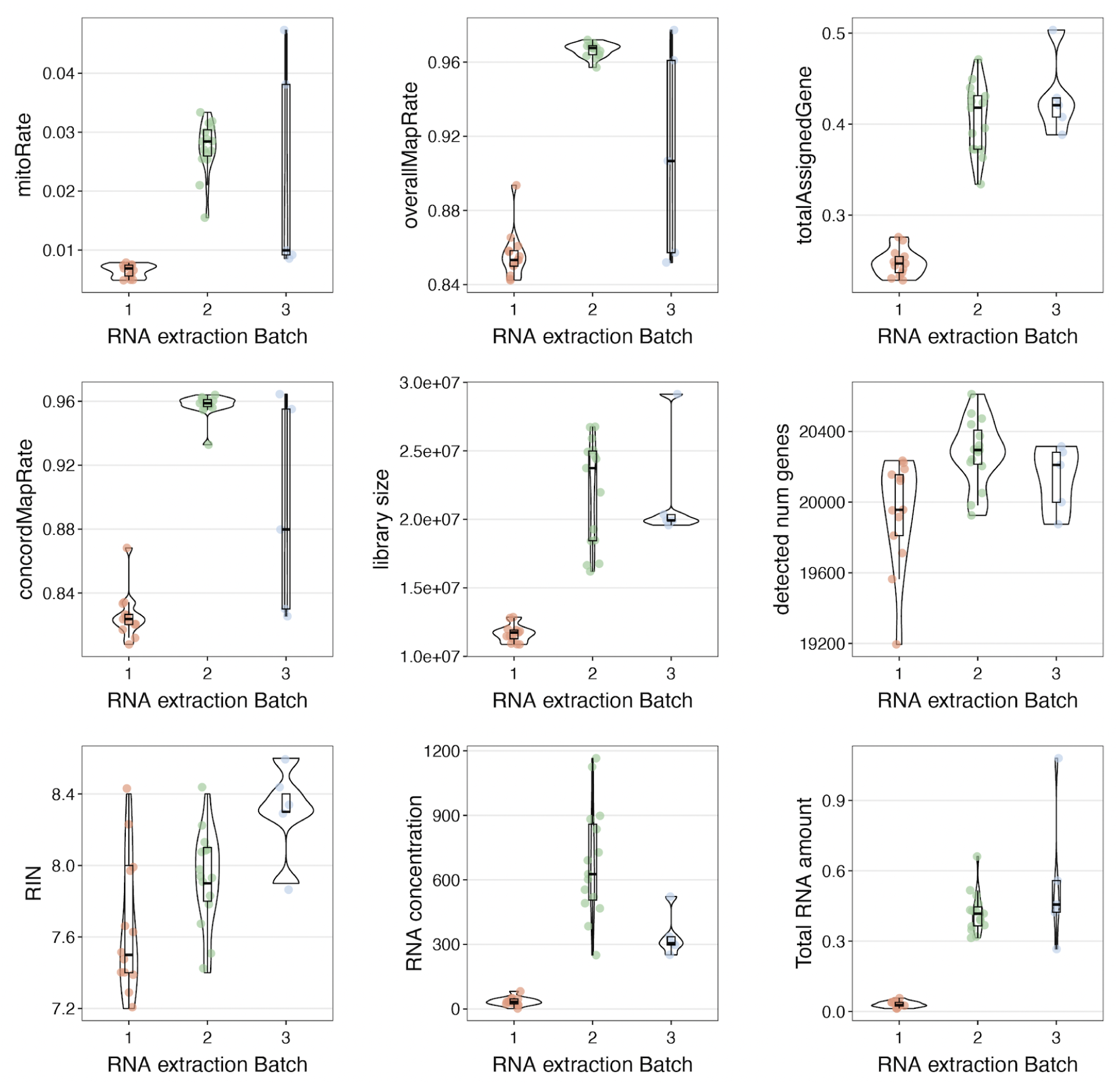
Quality control metrics for samples across RNA extraction batches. Comparison of QC metrics of samples from the first (only Hb samples), second (only Amyg samples), and third batch for RNA extraction (additional Hb and Amyg samples). See **Table S3** for the description of these QC metrics.

**Figure S4:**
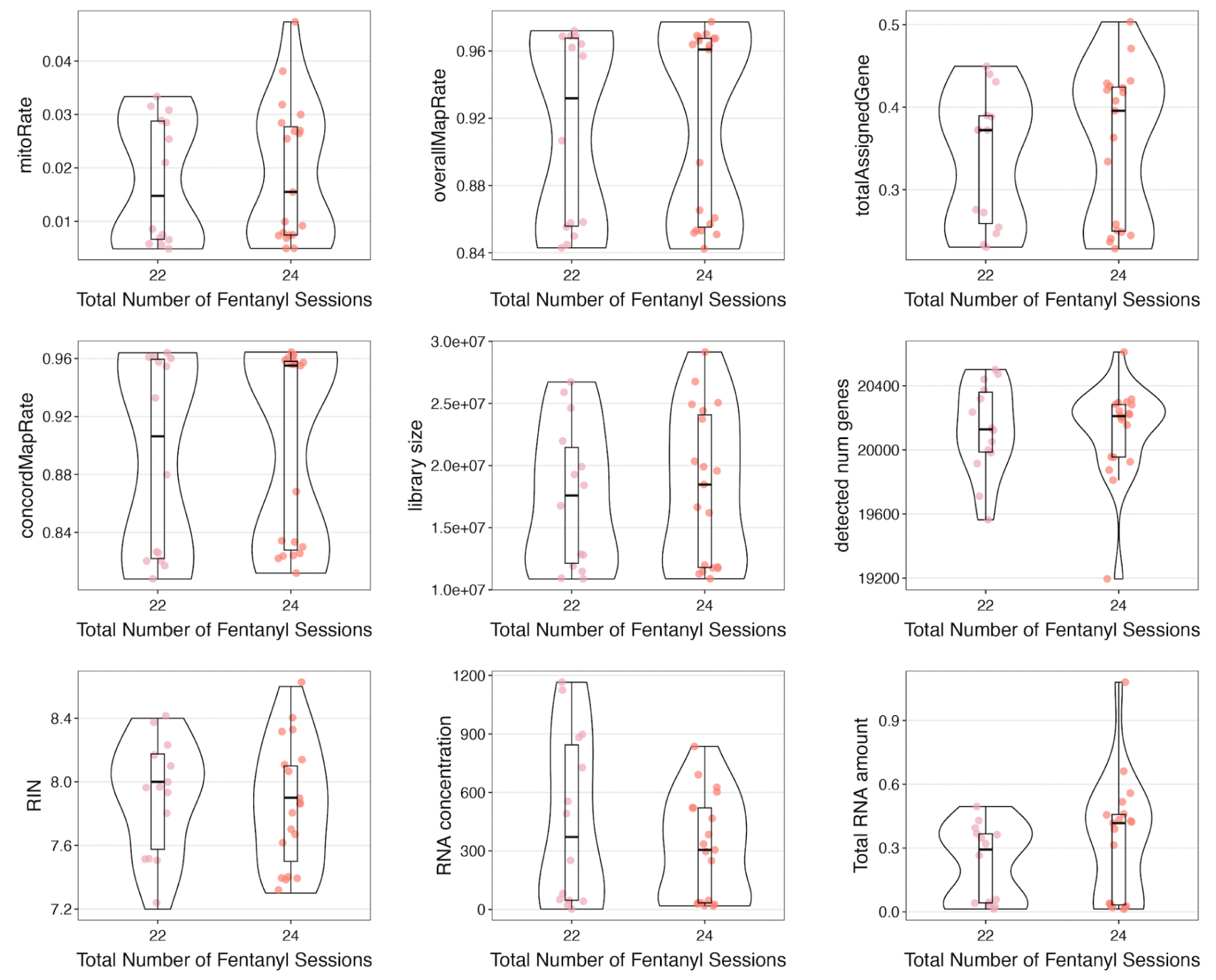
Quality control metrics for samples across total number of self-administration sessions. Comparison of QC metrics for (Hb and Amyg) samples from rats who had 22 and 24 total (fentanyl or saline) self-administration sessions. See **Table S3** for the description of these QC metrics.

**Figure S5:**
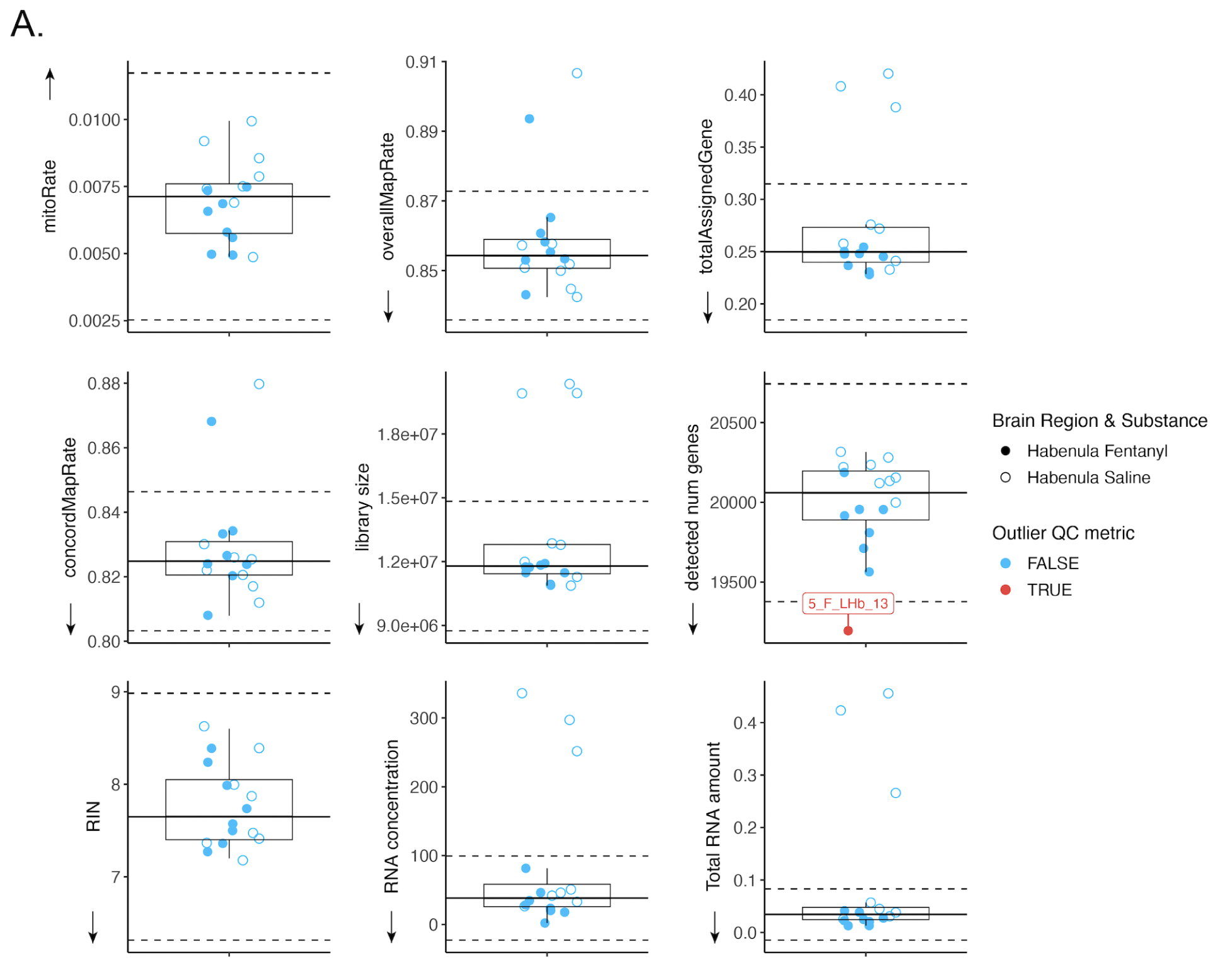
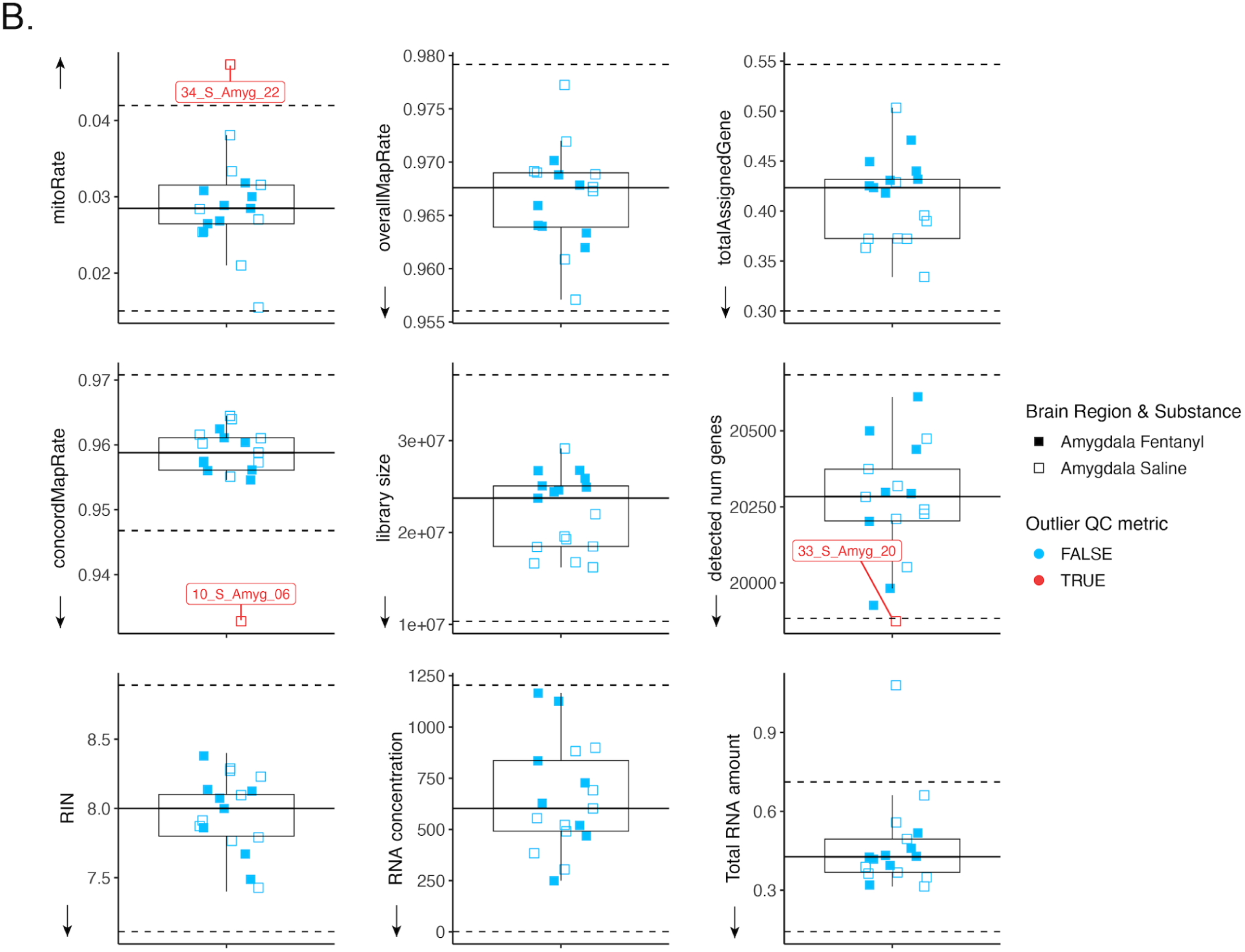
Low-quality sample identification. Detection of low-quality metrics for (**A**) Hb and (**B**) Amyg fentanyl (filled circles/squares) and saline (empty circles/squares) samples. QC metric outliers (in red) were identified as those being 3 median-absolute-deviations (MAD; dotted lines) away from the median (solid line). Only lower outliers were considered poor-quality for all QC metrics except `mitoRate`, for which higher outliers were considered instead (indicated by arrows). Samples with outlier QC metrics are labeled and were subjected to further evaluation in downstream analyses (**Figure S7**). See **Table S3** for the description of these QC metrics.

**Figure S6:**
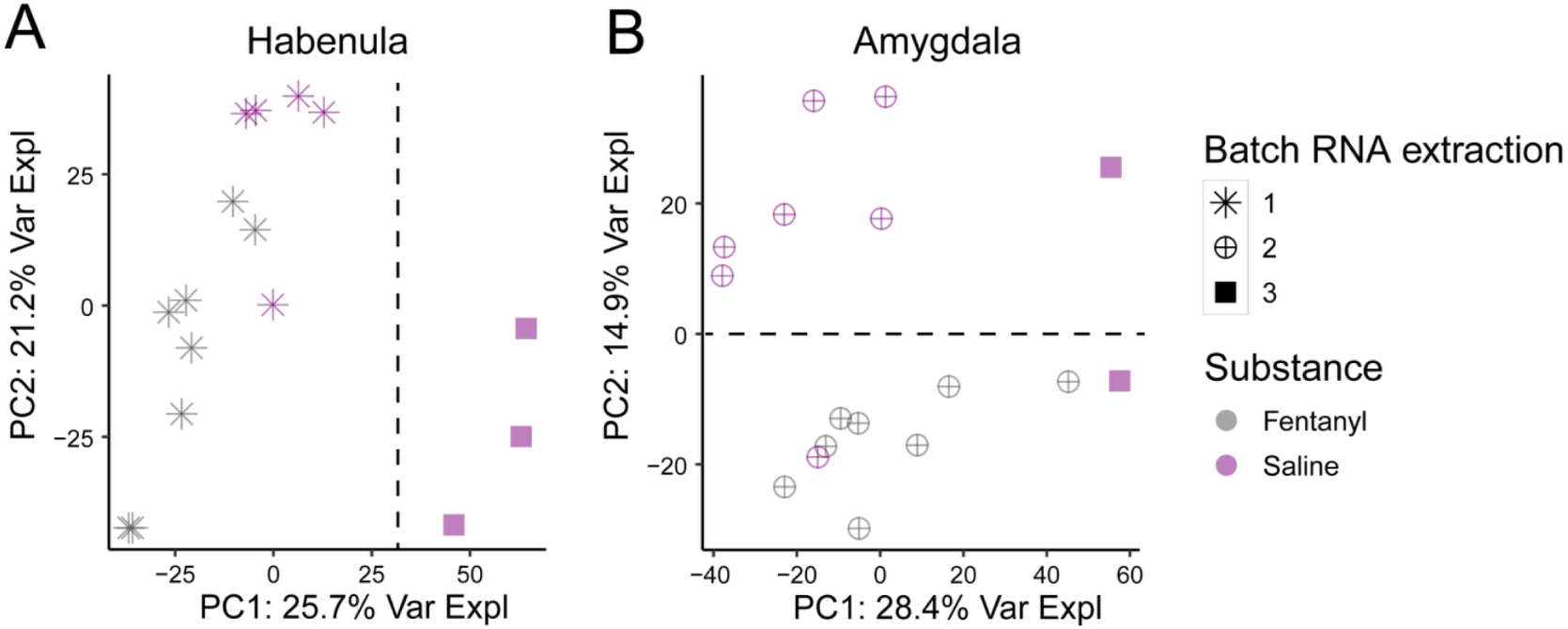
Principal Component Analysis. PC1 vs. PC2 for gene expression in (**A**) Hb and (**B**) Amyg samples. Percentages of variance explained by each PC are indicated on the axes. Samples are shaped by RNA extraction batch and colored by substance.

**Figure S7:**
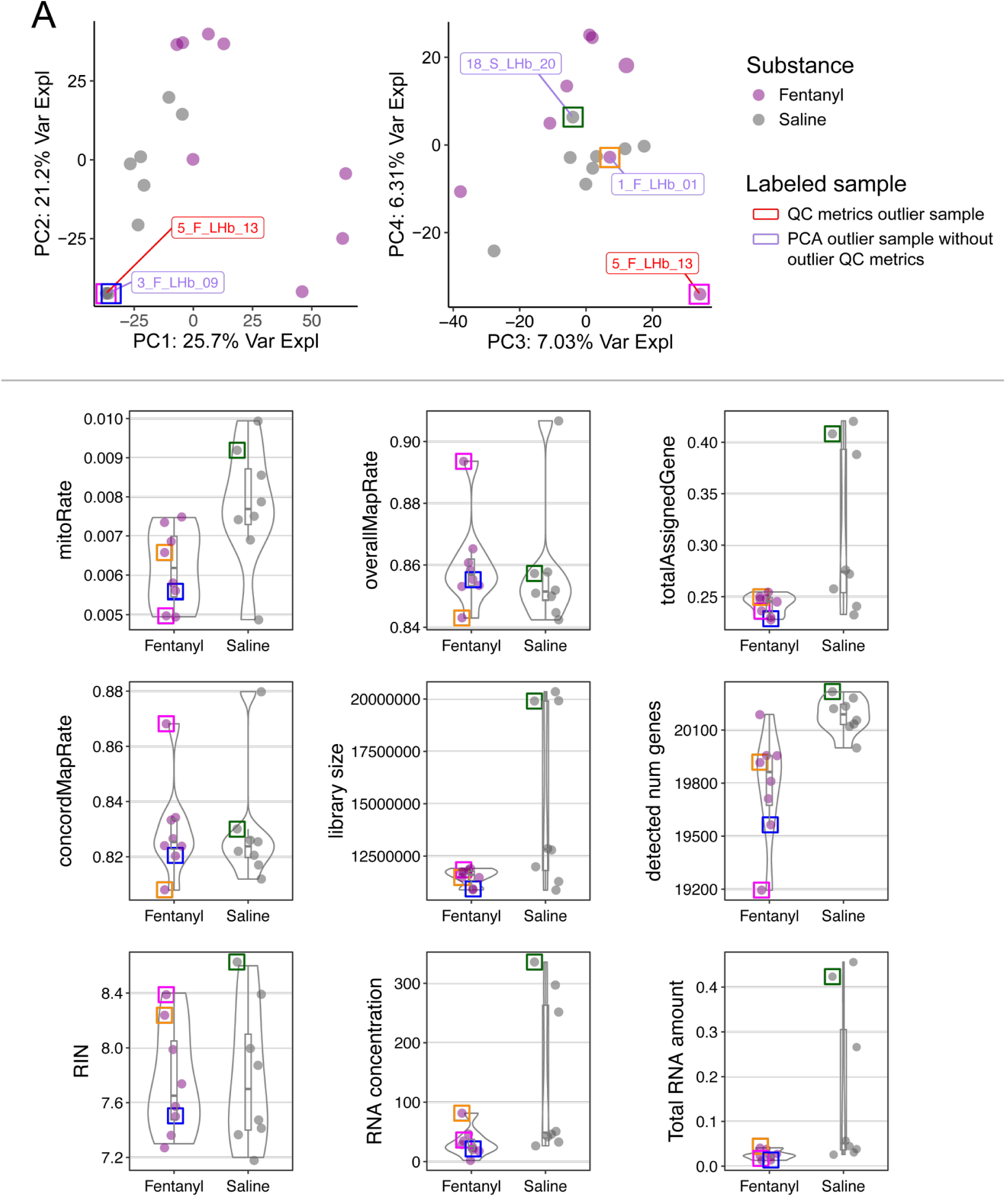
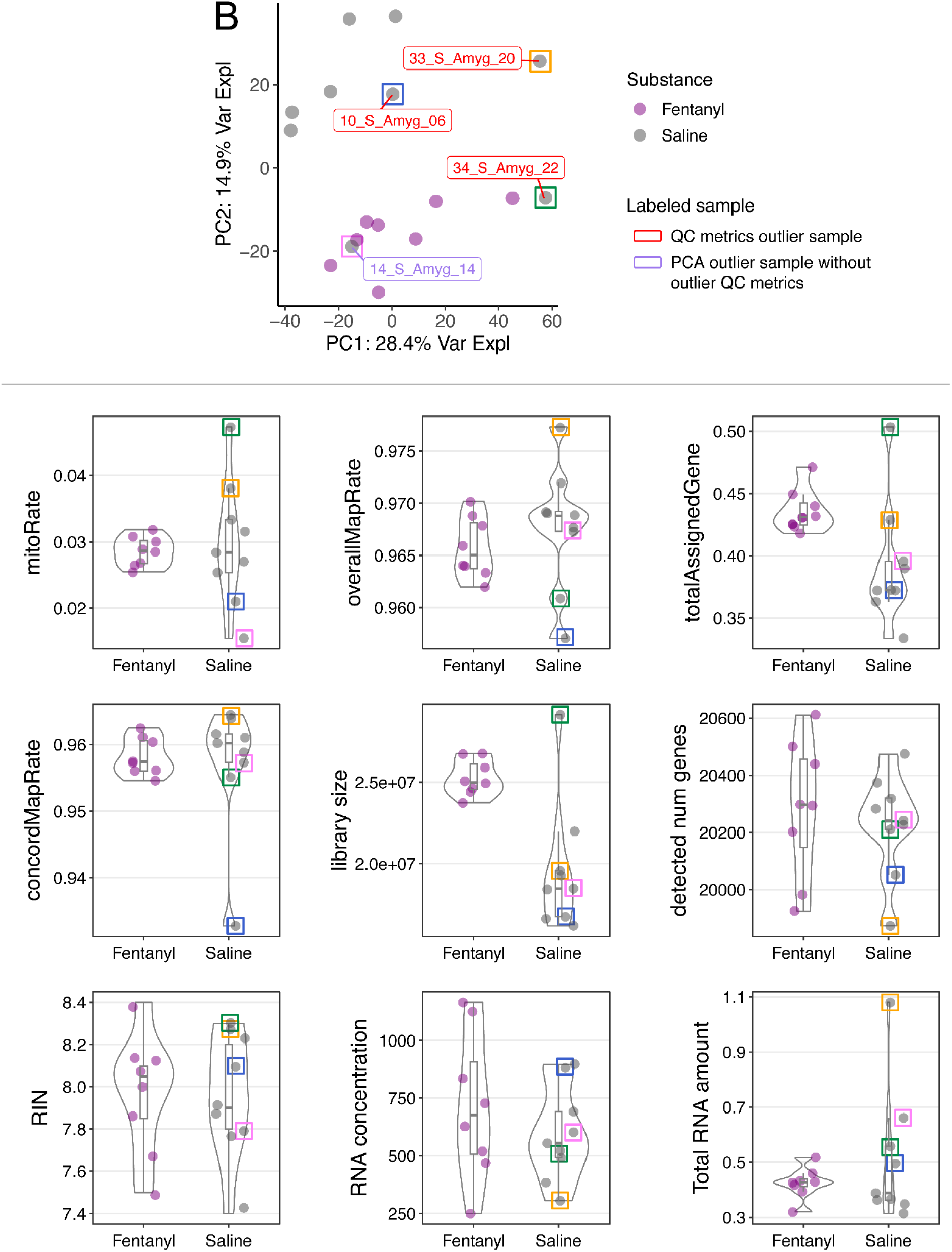
Manual sample quality examination based on PCA. PCx vs. PCy (top) for (**A**) Hb and (**B**) Amyg samples. QC metrics outlier samples are labeled in red (see **Figure S5**); samples segregated from the rest in each PC plot, as well as fentanyl and saline samples closer to samples from the other substance group were considered PCA outlier samples and are labeled in purple. The percentage of variance explained by each PC is shown on axis labels. For both, QC metrics and PCA outlier samples, all their QC metrics were reexamined (bottom box plots); different colored squares indicate the different outlier samples. Samples in all plots are colored by substance. See **Table S3** for the description of these QC metrics.

**Figure S8:**
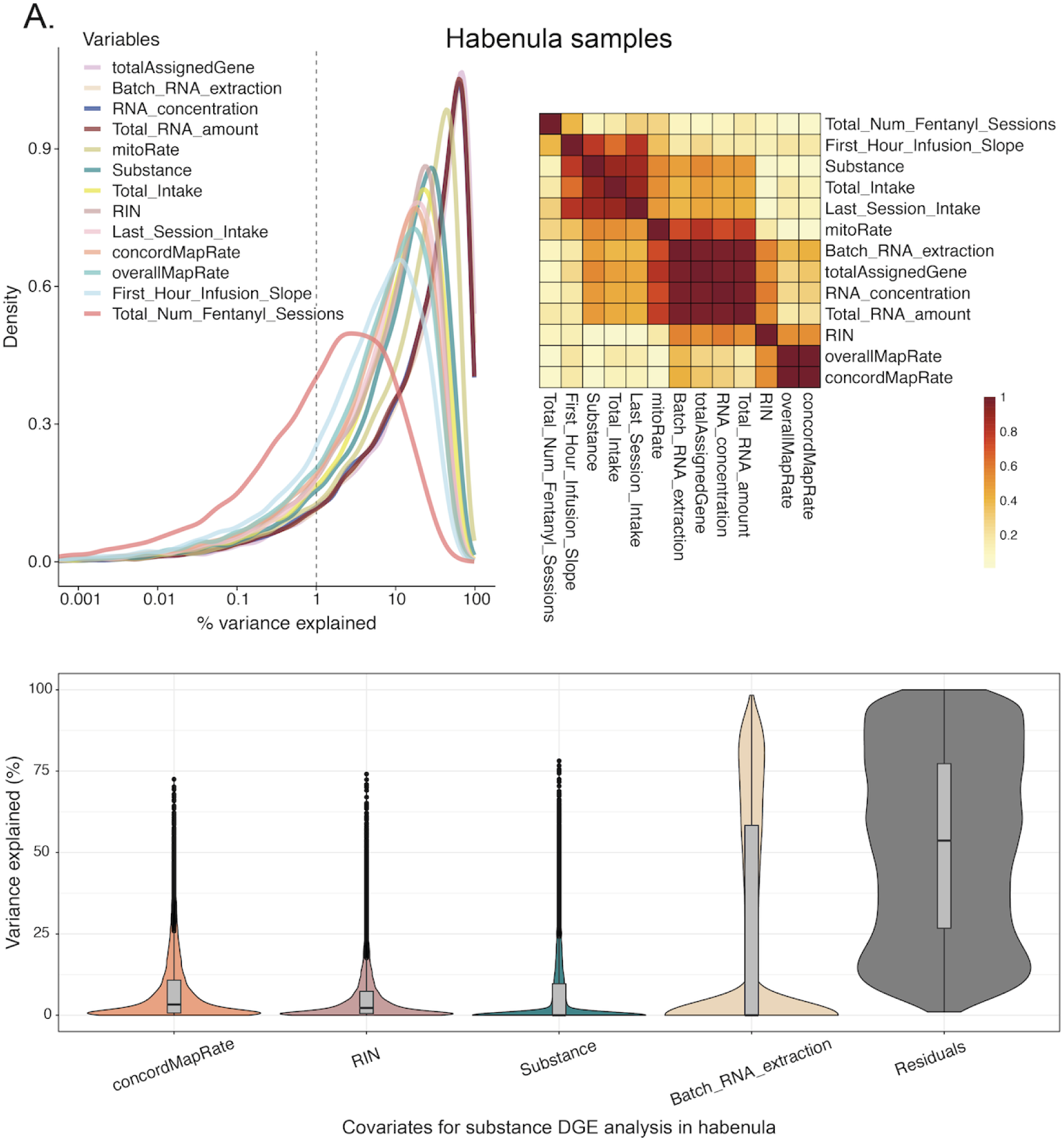
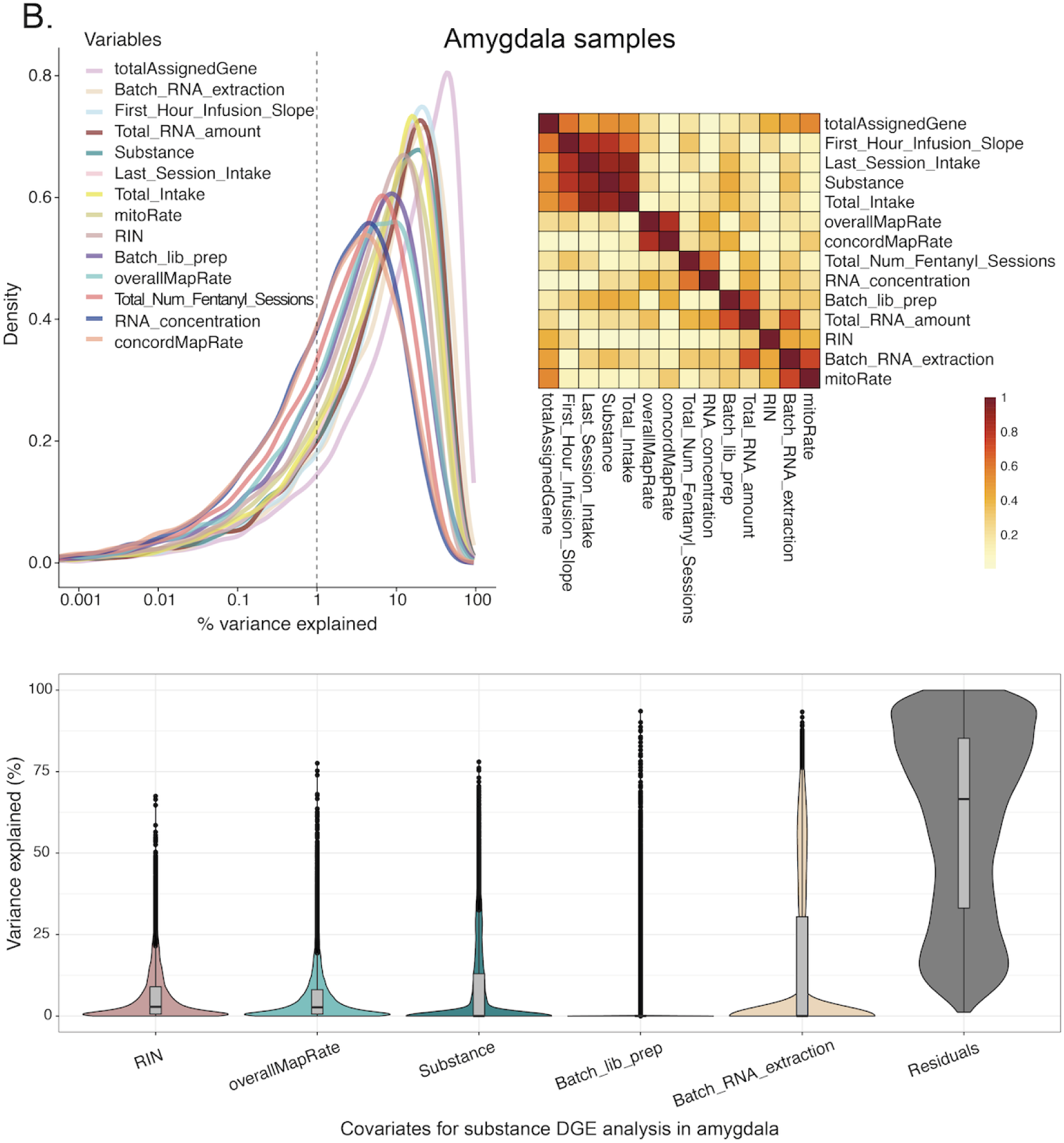
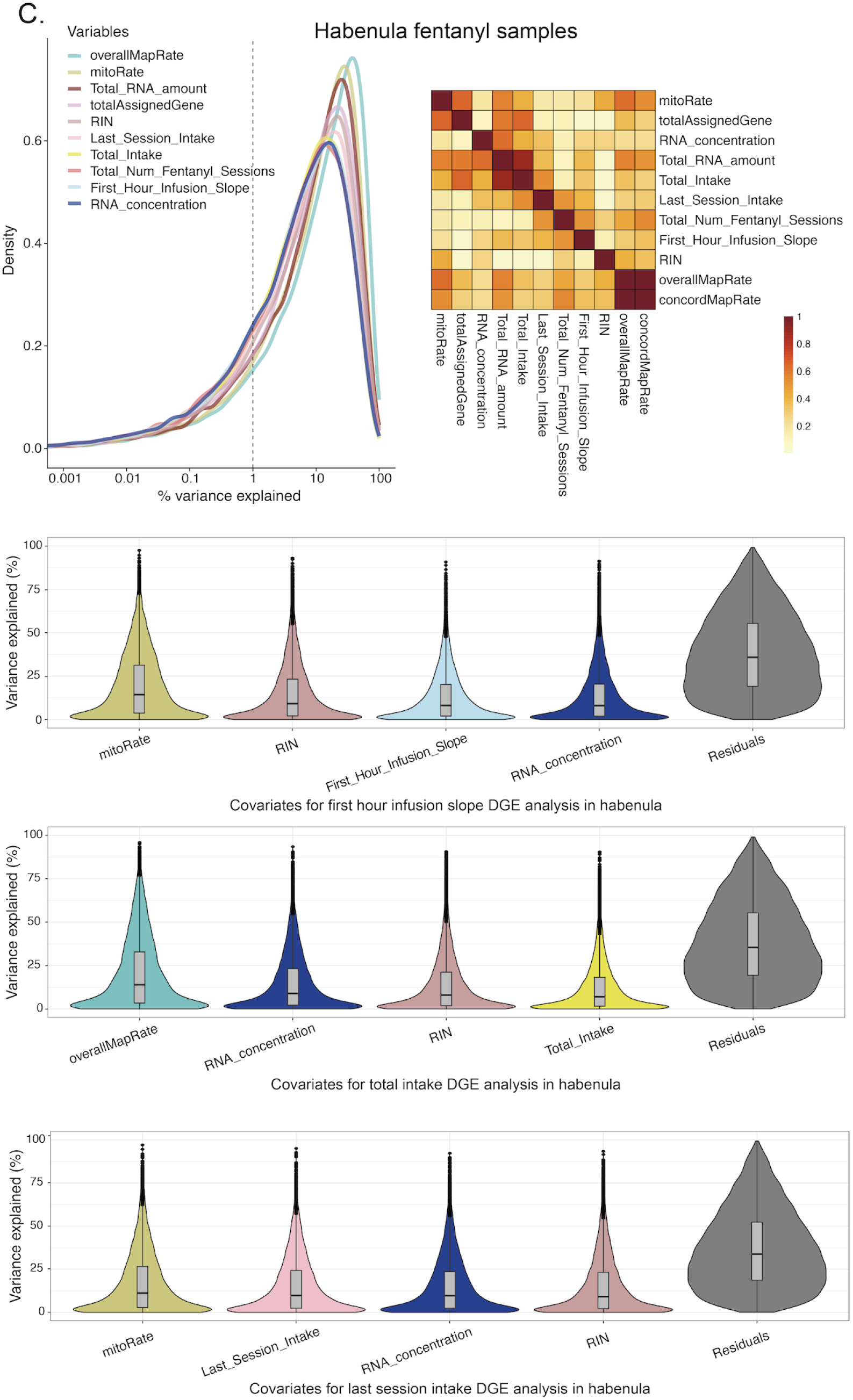
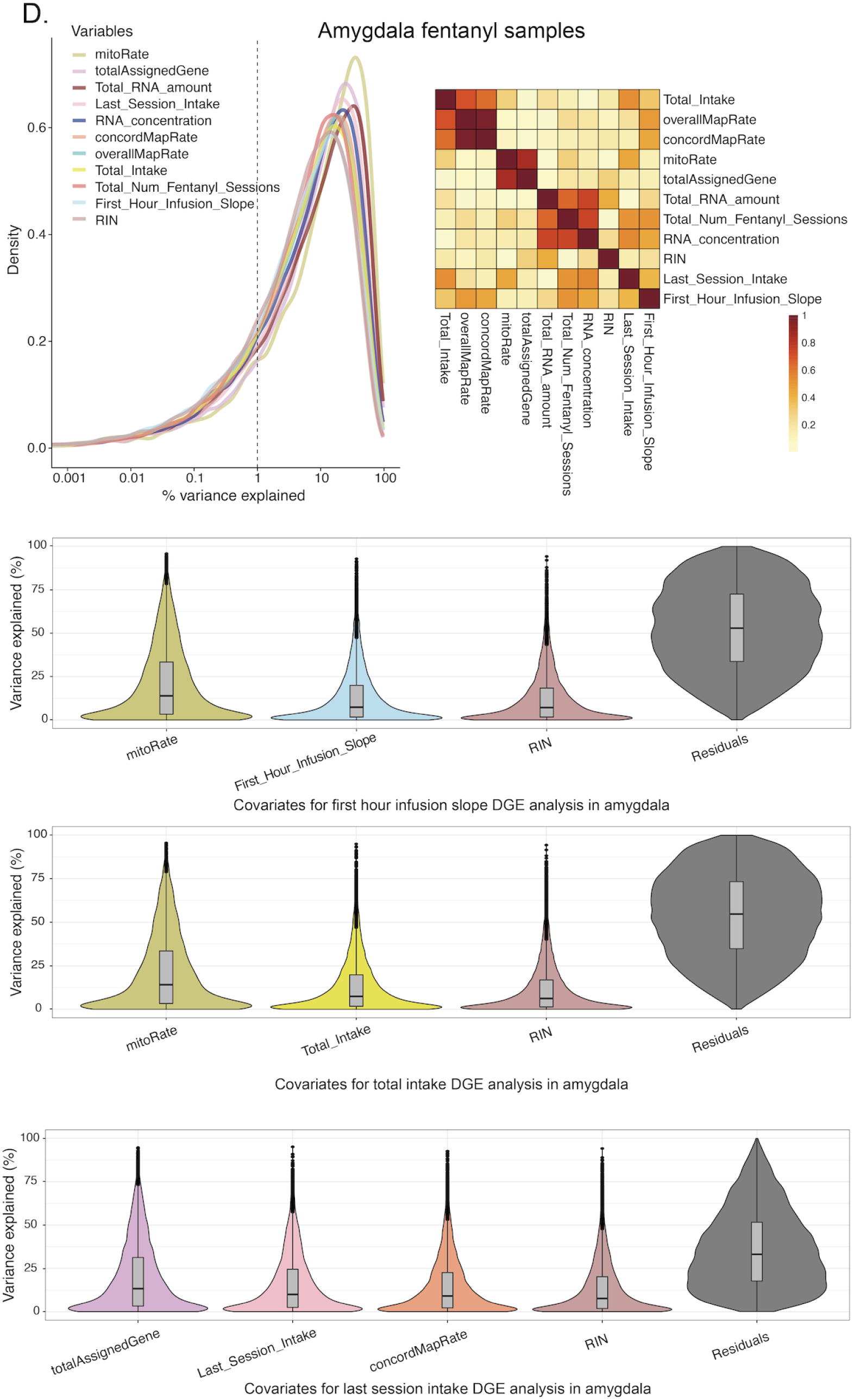
Sample-level covariate selection for DGE analysis. Gene expression variance partition analysis in **A.** Hb (all rats), **B.** Amyg (all rats), **C.** Hb (fentanyl rats only), and **D.** Amyg (fentanyl rats only). Top left: density plot for the percentages of variance explained in the expression of each gene by each sample-level variable. Top right: canonical correlation between each pair of variables. Variables included in the models for DGE analyses (A-B. for substance, and C-D. for rat behavioral traits) were selected based on their contributions to gene expression variance and correlations with other variables. Bottom: percentage of variance in the expression of each gene explained by each variable included in the DGE model, considering all other included variables in the model (x-axis); variables are ordered by decreasing median percentage of variance explained. Related to **Figure 2**. See **Table S3** for the description of these variables and QC metrics.

**Figure S9:**
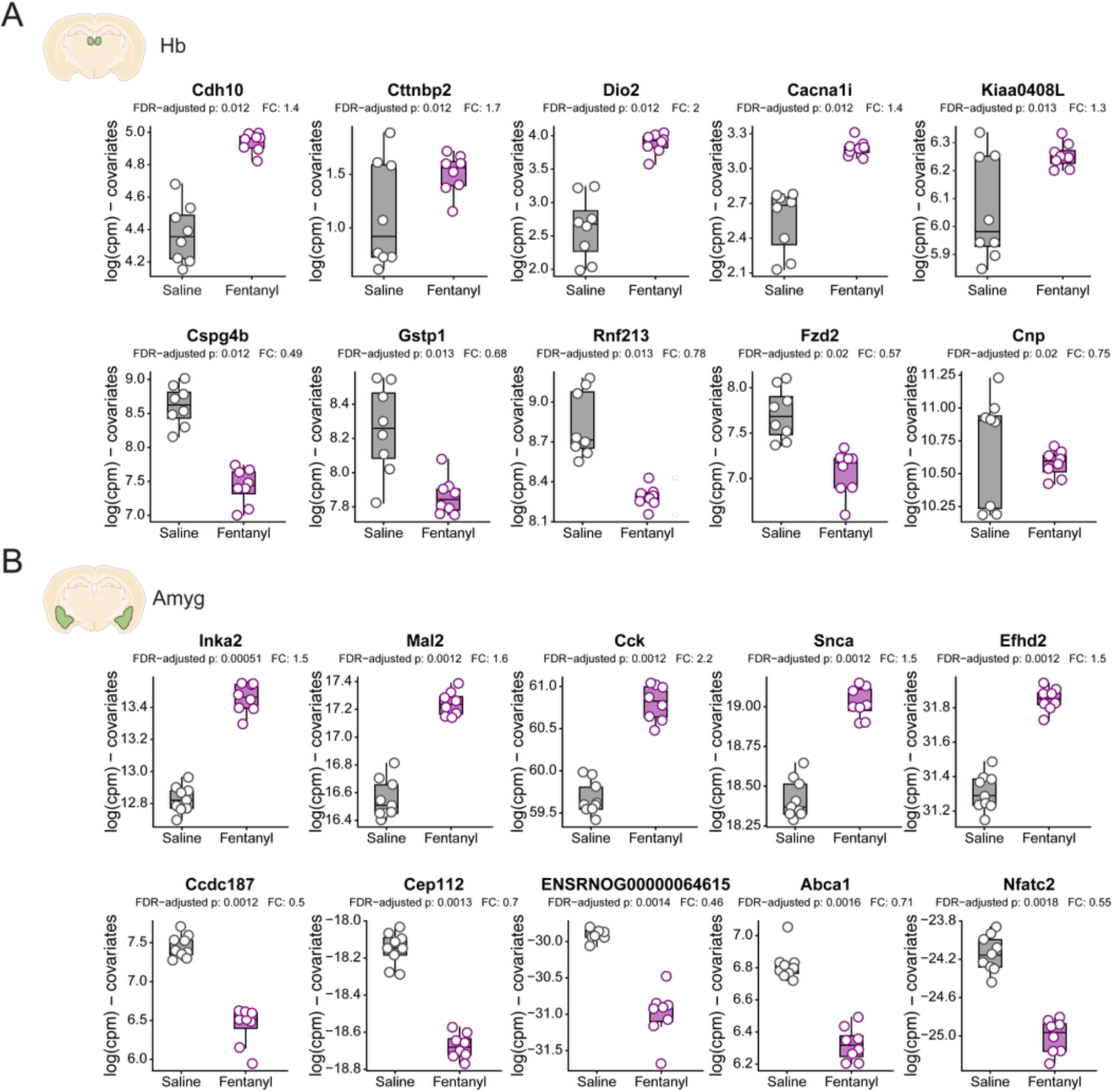
Top 5 differentially expressed genes in Hb and Amyg following chronic LgA fentanyl self-administration. (**A-B**) Box plots showing the expression of the top five up- and down-regulated DEGs for fentanyl vs. saline in Hb (**A**) and Amyg (**B**). Gene expression is given in log_2_(CPM) after regressing out covariates. FDR-adjusted p-value and fold change (FC) are shown for each gene. Boxes extend from the 25^th^ to 75^th^ percentiles; lines within the boxes represent the median; whiskers indicate the minimum and maximum values, superimposed with individual rat data points. Fentanyl Hb n = 8; Saline Hb n = 8; Fentanyl Amyg n = 8; Saline Amyg n = 9. Related to **Figure 2**.

**Figure S10:**
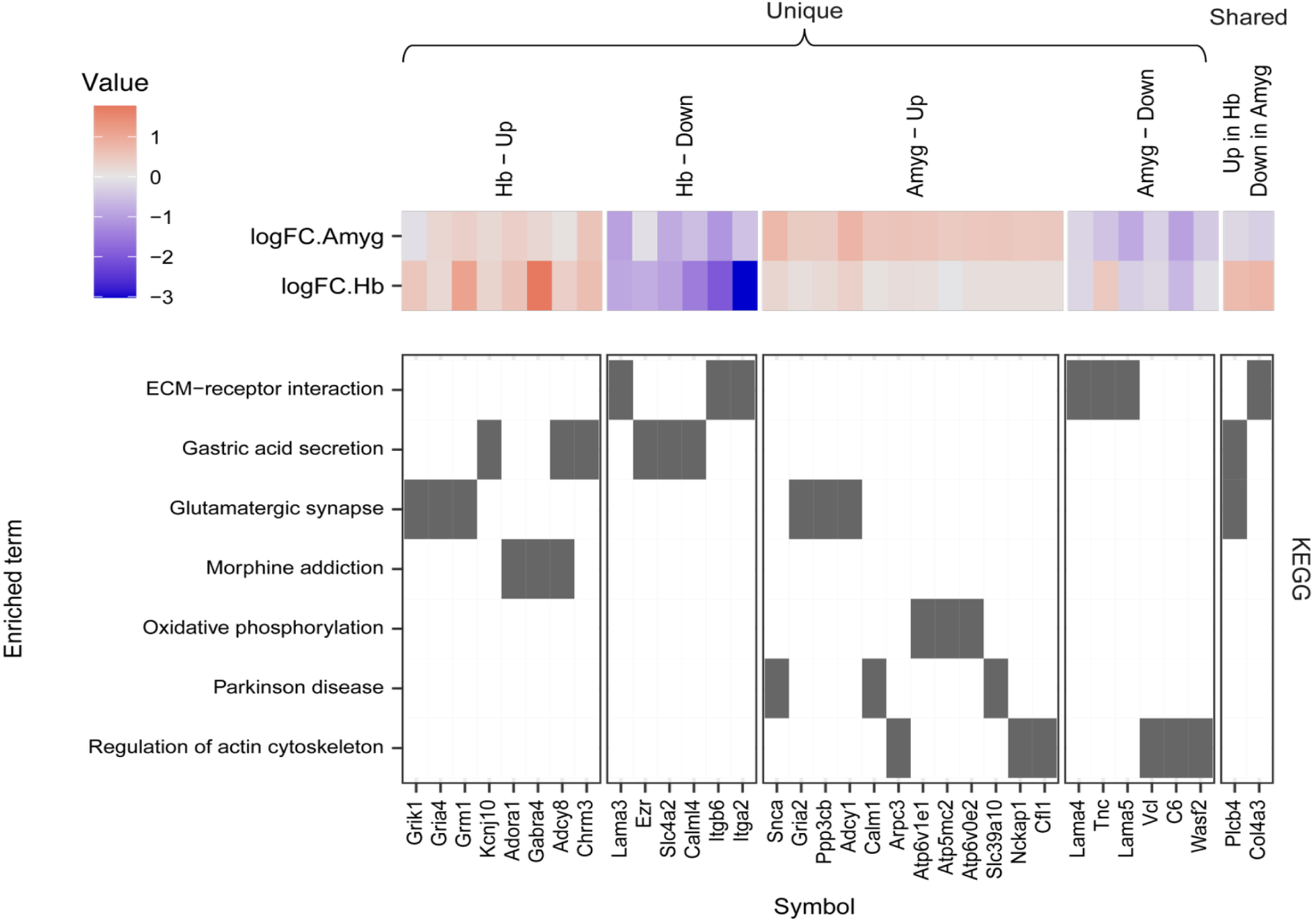
Biological KEGG pathways dysregulated by chronic fentanyl self-administration in Hb and Amyg. Tile plot displays DEG (x-axis) membership to an enriched pathway as a filled tile. Key DEGs from each pathway are shown, categorized by their unique or shared up- and down-regulation in Hb and Amyg. Top heatmap shows DEG mean-centered log_2_FC in Hb and Amyg. Related to **Figure 2**, **Table S8**, **Table S9**.

**Figure S11:**
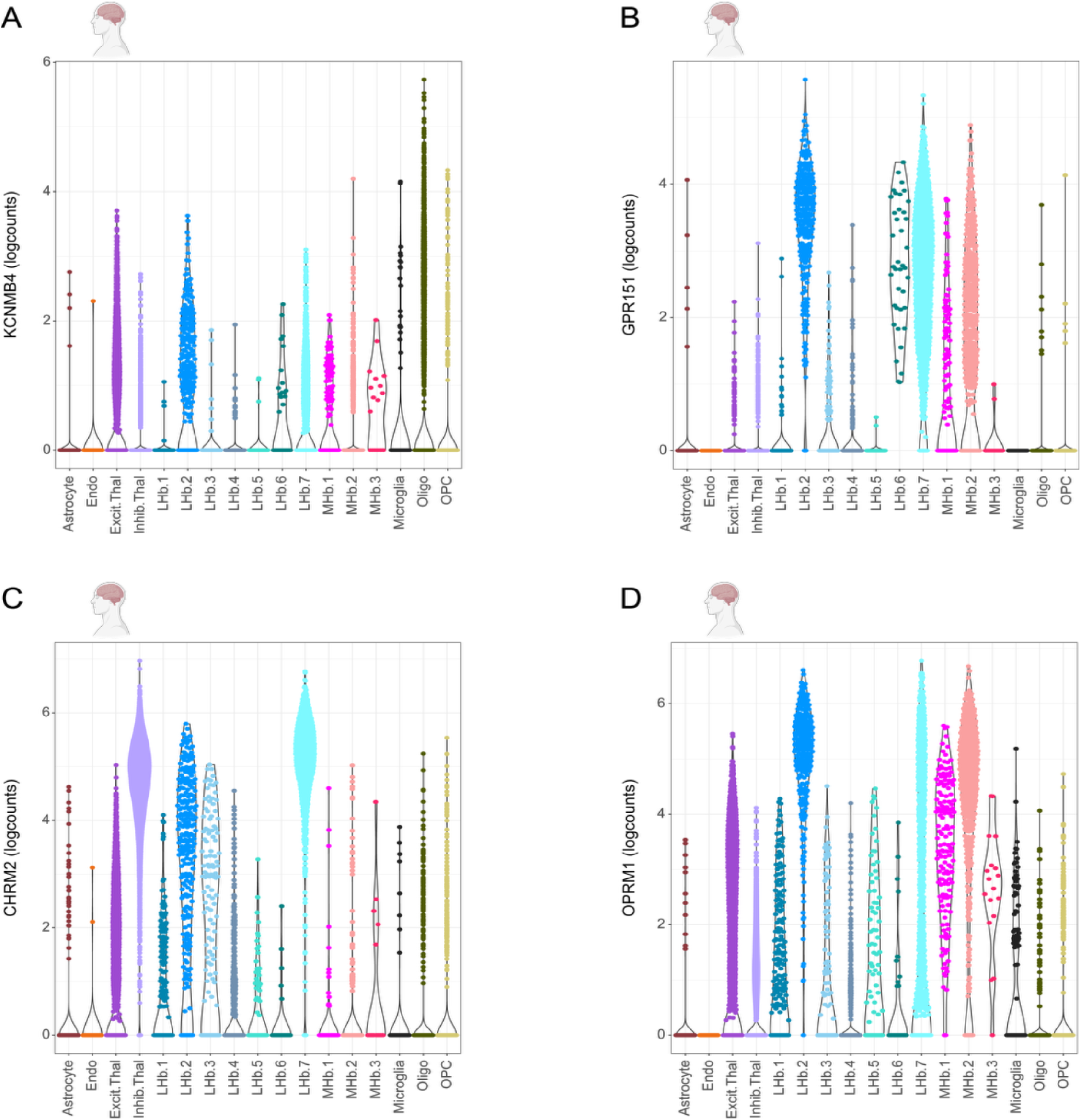
Expression of *KCNMB4*, *GPR151, CHRM2* and *OPRM1* in human habenula cell types. (A-D) Violin plots showing expression of (**A**) *KCNMB4*, (**B**) *GPR151*, (**C**) *CHRM2*, and (**D**) *OPRM1* in human Hb cell types from Yalcinbas et al., 2025 [52]. *Kcnmb4*, *Gpr151*, and *Chrm2* mark the mouse LHb.6 subpopulation identified by Hashikawa et. al. [51], which we found to be enriched in our rat upregulated Hb fentanyl DEGs. These genes are highly expressed in human LHb.2 and LHb.7 subpopulations, which also express *OPRM1*. This suggests that fentanyl-sensitive mouse LHb.6 may be conserved with these *OPRM1*-expressing human LHb.2 and LHb.7 neuronal populations. Related to **Figure 3**.

## Supplementary Tables

**Table S1: Behavioral raw data per LgA self-administration session per rat.** Session-by-session raw behavioral data pertaining to operant (lever press) behaviors and infusions from each rat across all long-access self-administration sessions. See **Table S3** for the description of these variables.

**Table S2: Behavioral rat data in LgA sessions.** Individual rat data for behavioral covariates relating to fentanyl intake and intake escalation across long-access self-administration sessions. See **Table S3** for the description of these variables.

**Table S3: Dictionary of sample variables.** Description of sample/rat variables analyzed throughout the study. Related to **Table S1**, **Table S2**, and **Table S4**.

**Table S4: Sample metadata and QC metrics.** Sample level variables analyzed, including data regarding rat self-administration sessions and sample batches for RNA extraction, library preparation, and sequencing, as well as quality control metrics. See **Table S3** for the description of these variables.

**Table S5: DEGs for substance in Hb.** Metadata, *limma* DE statistics, and Ensembl gene annotation for DEGs obtained for fentanyl vs. saline in Hb. See *limma* [65] documentation for these statistics definitions. Related to **Figure 2, Table S8**, **Table S10**.

**Table S6: DEGs for substance in Amyg.** Metadata, *limma* DE statistics, and Ensembl gene annotation for DEGs obtained for fentanyl vs. saline in Amyg. See *limma* [65] documentation for these statistics definitions. Related to **Figure 2, Table S9, Table S10**.

**Table S7. Common DEGs for substance in habenula and amygdala.** Metadata, region-specific *limma* DE statistics, and Ensembl gene annotation for overlapping DEGs for fentanyl vs. saline in habenula and amygdala. See *limma* [65] documentation for these statistics definitions. Related to **Figure 2, Table S5, Table S6**.

Table S8. Functional enrichment results for substance DEGs in Hb. GO terms for biological processes (BP), molecular functions (MF), cellular components (CC), and KEGG pathways that are significantly enriched in up- and down-regulated DEGs for fentanyl vs. saline in Hb. Provided are the ID and description of each significant term/pathway, the number and fraction of up/down-regulated DEGs annotated to each term (`Count` and `GeneRatio`, respectively), as well as the list of such genes (`geneID`), the fraction of genes in universe annotated to each term (`BgRatio`), fold of enrichment, *p*-value, and FDR-corrected *p*-value. Related to **Figure 2**, **Figure S10**, **Table S5**.

**Table S9. Functional enrichment results for substance DEGs in Amyg.** Same as **Table S8** but for up- and down-regulated DEGs for fentanyl vs. saline in Amyg. Related to **Figure 2**, **Figure S10**, **Table S6**.

**Table S10: Results for all DGE analyses and genes in Hb and Amyg.** Gene-level metadata and *limma* DE statistics of each gene for substance and rat behavior DGE analyses (fentanyl vs. saline, first hour infusion slope, total intake, and last session intake) in Hb and Amyg. See *limma* [65] documentation for these statistics definitions. Related to **Figure 2**. This table includes all the data from **Table S5**, **Table S6**, and **Table S7**.

**Table S11: Top 100 Mea*nRatio* marker genes per cell type in mouse Hb.** For all cell subpopulations and Hb neuronal subpopulations in the Hb complex of control mice obtained in Hashikawa et al., 2020 [51], the top 100 most specific marker genes for each based on the *MeanRatio* method, are reported. See *DeconvoBuddies* [67] documentation for column description. `Cell_type_resolution` column corresponds to the resolution of the cell subpopulation for which the gene is a marker. Related to **Figure 3**.

**Table S12: Top 50 *MeanRatio* marker genes per cell type in human Hb.** For broad and fine cell types in the human Hb-enriched epithalamus of neurotypical control donors obtained in Yalcinbas et al., 2025 [52], the top 50 most specific marker genes for each based on the *MeanRatio* method, are reported. See *DeconvoBuddies* [67] documentation for column description. `Cell_type_resolution` column corresponds to the resolution of the cell type for which the gene is a marker. Related to **Figure 3**.

**Table S13: Top 100 *MeanRatio* marker genes per cell type in rat Amyg.** For main cell types and inhibitory neuronal subtypes in the Amyg of control rats obtained in Zhou et al., 2023 [54], the top 100 most specific marker genes for each based on the *MeanRatio* method, are reported. See *DeconvoBuddies* [67] documentation for column description. `Cell_type_resolution` column corresponds to the resolution of the cell type for which the gene is a marker. Related to **Figure 4**.

**Table S14: Top 100 MeanRatio marker genes per cell type in human Amyg.** For broad and fine cell types in the human Amyg of neurotypical control donors obtained in Yu et al., 2023 [53], the top 100 most specific marker genes for each based on the *MeanRatio* method, are reported. See *DeconvoBuddies* [67] documentation for column description. `Cell_type_resolution` column corresponds to the resolution of the cell type for which the gene is a marker. Related to **Figure 4**.

## Notes

https://github.com/LieberInstitute/fentanyl_rat_hb_amy

https://www.ncbi.nlm.nih.gov/bioproject/PRJNA1179901

### Competing Interest Statement

The authors have declared no competing interest.

### Summary of Updates

In this revised version we have updated Figure 4D, Supplementary Figure 1E, expanded the main text, and added additional relevant references.

